# *Ex vivo* tissue perturbations coupled to single cell RNA-seq reveal multi-lineage cell circuit dynamics in human lung fibrogenesis

**DOI:** 10.1101/2023.01.16.524219

**Authors:** Niklas J. Lang, Janine Gote-Schniering, Diana Porras-Gonzalez, Lin Yang, Laurens J. De Sadeleer, R. Christoph Jentzsch, Vladimir A. Shitov, Shuhong Zhou, Meshal Ansari, Ahmed Agami, Christoph H. Mayr, Baharak Hooshiar Kashani, Yuexin Chen, Lukas Heumos, Jeanine C. Pestoni, Emiel Geeraerts, Vincent Anquetil, Laurent Saniere, Melanie Wögrath, Michael Gerckens, Rudolf Hatz, Nikolaus Kneidinger, Jürgen Behr, Wim A. Wuyts, Mircea-Gabriel Stoleriu, Malte D. Luecken, Fabian J. Theis, Gerald Burgstaller, Herbert B. Schiller

**Author notes:** … these authors contributed equally (Niklas J. Lang, Janine Gote-Schniering).

## Abstract

Pulmonary fibrosis develops as a consequence of failed regeneration after injury. Analyzing mechanisms of regeneration and fibrogenesis directly in human tissue has been hampered by the lack of organotypic models and analytical techniques. In this work, we coupled *ex vivo* cytokine and drug perturbations of human precision-cut lung slices (hPCLS) with scRNAseq and induced a multi-lineage circuit of fibrogenic cell states in hPCLS, which we show to be highly similar to the *in vivo* cell circuit in a multi-cohort lung cell atlas from pulmonary fibrosis patients. Using micro-CT staged patient tissues, we characterized the appearance and interaction of myofibroblasts, an ectopic endothelial cell state and basaloid epithelial cells in the thickened alveolar septum of early-stage lung fibrosis. Induction of these states in the *ex vivo* hPCLS model provides evidence that the basaloid cell state was derived from alveolar type-2 cells, whereas the ectopic endothelial cell state emerged from capillary cell plasticity. Cell-cell communication routes in patients were largely conserved in the hPCLS model and anti-fibrotic drug treatments showed highly cell type specific effects. Our work provides an experimental framework for perturbational single cell genomics directly in human lung tissue that enables analysis of tissue homeostasis, regeneration and pathology. We further demonstrate that hPCLS offers novel avenues for scalable, high-resolution drug testing to accelerate anti-fibrotic drug development and translation.

## INTRODUCTION

Fibrogenesis, the activation of stromal cells to produce an excess of extracellular matrix (ECM), is a central pillar of physiological tissue repair that is dysregulated in various diseases and causes significant morbidity and mortality, accounting for 45% of deaths in the developed world^1^. Pulmonary fibrosis (PF), the result of progressing and non-resolving fibrogenesis due to failed lung regeneration, affects more than 500,000 patients in Europe^2^. Rather than a disease itself, progressing PF is regarded as a pathobiological mechanism present in multiple entities of interstitial lung diseases, causing excessive scarring and architectural distortion of the lung tissue, ultimately resulting in respiratory failure and death in most patients. The median survival without treatment in idiopathic pulmonary fibrosis (IPF), the most prevalent form of PF, is only 3-5 years^2^. Two drugs, Nintedanib and Pirfenidone, are approved for clinical use in PF^3–5^; however, both only slow down disease progression and do not restore normal tissue architecture^6, 7^.

Recent single cell RNA sequencing (scRNA-seq) studies of PF patient cohorts^8–12^ have led to the molecular description of fibrosis-specific cell states, including the KR17+/KRT5-basaloid epithelial cells, as well as several novel disease-associated endothelial, stromal and immune cell states. Many of these cell states described in PF also appeared in the lungs during severe COVID-19 and consecutive acute respiratory distress syndrome^13–17^. This suggests that many PF-associated cell states may represent Regenerative Intermediate Cell States (RICS) that appear after lung injury and potentially persist in chronic disease.

In lung regeneration, the fibrogenic phase is transient, and the ECM produced by fibroblasts is removed upon completion of repair. We have previously used longitudinal single cell RNA-seq to chart the evolution of RICS during mouse lung fibrogenesis and regeneration^18, 19^. Lung injury induced the transient appearance of distinct epithelial, endothelial, stromal, and immune RICS, which may form interconnected cellular circuits that orchestrate fibrogenesis during regeneration and become distorted during disease. RICS are limited in time and space by mostly unknown regulatory mechanisms. The spatiotemporal and functional interdependencies of RICS during human lung fibrogenesis, which would provide a conceptual framework for discovering novel drug targets for PF, are currently unknown.

The lack of translatable mouse models compromises the functional and spatiotemporal characterization of disease-relevant RICS. As a consequence, current drug discovery efforts primarily focus on the modulation of specific molecular pathways in highly artificial conditions using cell lines or, more recently, organoid and co-culture assays, which mostly fail to recapitulate the tissue niches in which a drug candidate has to unfold its anti-fibrotic effect in the patient. Hence, therapeutically relevant cell-cell communication can only be poorly modeled using such *in vitro* cell culture systems. Human precision-cut lung slices (hPCLS) have emerged as a promising new *ex vivo* model to mechanistically study lung fibrogenesis^20–22^ . hPCLS have the unique advantage of retaining the full cellular diversity in its native 3D architecture, thereby allowing functional characterization of RICS within their preserved tissue niches and cell-cell communication circuits.

In this work, we use single cell RNA-seq to analyze the cytokine- and drug-induced perturbation response of cells in hPCLS. We benchmark the hPCLS perturbation data against *in vivo* scRNA-seq data from PF patients, revealing the strengths and limitations of the *ex vivo* hPCLS model. We demonstrate *ex vivo* induction of RICS in hPCLS with highly similar transcriptional profiles as their *in vivo* counterparts in PF and validate the appearance of these states in early-stage disease using immunofluorescence analysis of micro-CT staged tissues from PF patients. Finally, we analyze drug effects on multi-lineage circuits of RICS, identifying established and novel drug-induced cell state changes and drug-specific cellular communication patterns. Using deep-learning-based query-to-reference mapping, we illustrate proof-of-concept phenotypic drug mode of action results, establishing a workflow for scalable testing of new anti-fibrotic drugs and their combinations in hPCLS.

## RESULTS

### Perturbation analysis in human lung tissue at single cell resolution

We generated hPCLS from non-fibrotic human lung tissues and treated them with a cocktail of pro-fibrotic cytokines consisting of TGF-β, PDGF-AB, TNF-α, and LPA (FC)^20, 23^, or a control cocktail of solvents used for the cytokine mix (CC) (**Fig 1a**). To analyze the effects of anti-fibrotic drugs, we co-treated FC-stimulated hPCLS with the clinically approved anti-fibrotic drug Nintedanib^3^ (FC+Nintedanib), as well as an *N*-(2-butoxyphenyl)-3-(phenyl)acrylamide (N23P) derivative of Tranilast (CMP4), which we recently have identified as a novel anti-fibrotic drug candidate (FC+CMP4)^24^. After six days, we performed single-cell RNA sequencing (scRNA-seq) using the 10x Genomics Chromium platform (**Fig. 1a**). In total, we profiled 63,581 single-cells from two tissue donors (**Supp. Fig. 1a - g**) across four treatment conditions (**Fig. 1b and c**).

**Figure 1.**
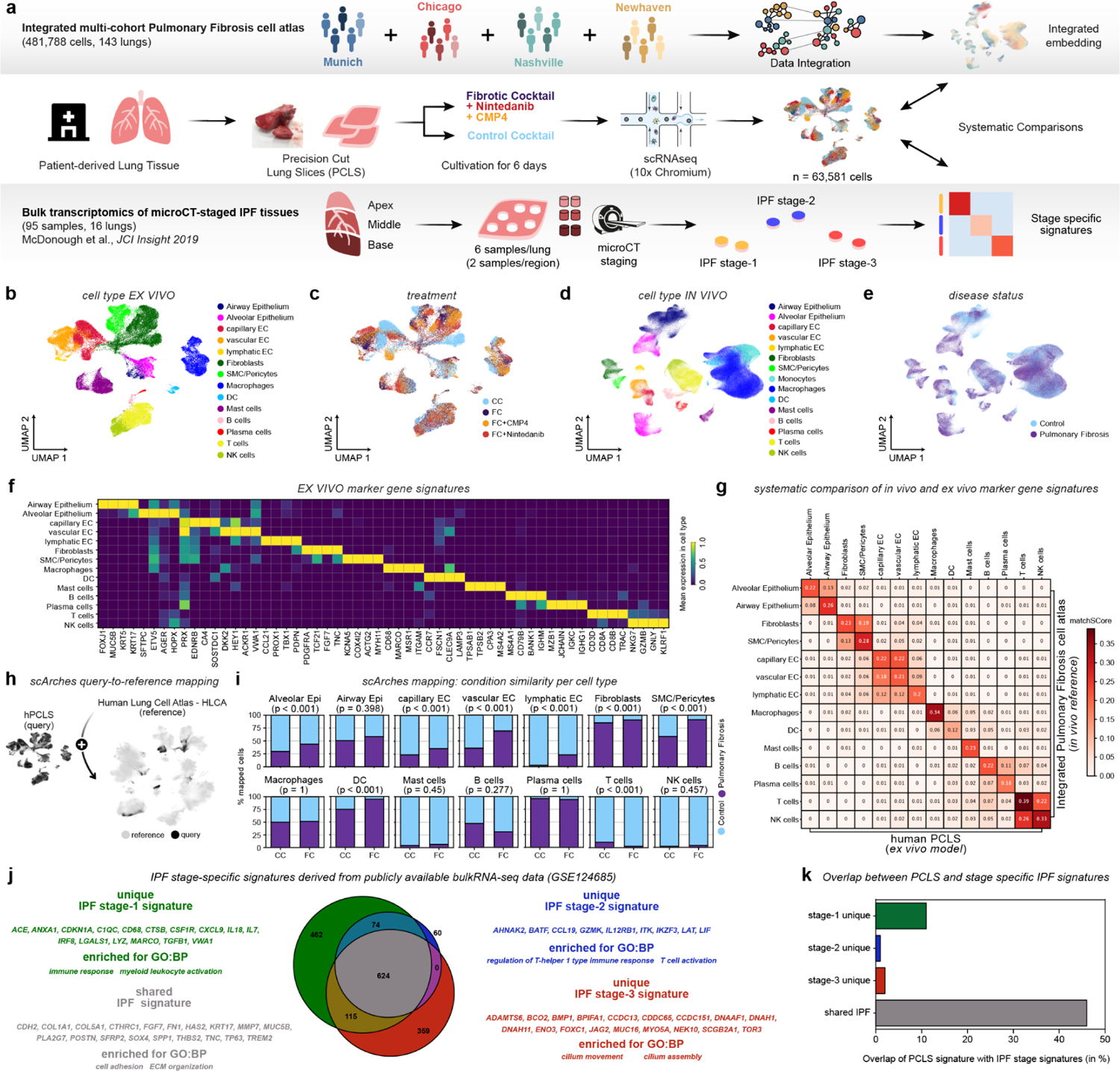
An *ex vivo* model of human lung fibrogenesis recapitulates early-stage events in PF patients. **(a)** Benchmarking strategy: Human precision-cut lung slices (hPCLS) were generated from uninvolved peritumoral tissue and treated under the indicated conditions before single cell RNA-seq analysis (FC = Fibrotic Cocktail: TGFb1, PDGFb, TNFa, and LPA, CC = Control Cocktail: diluents of the components) at day six of treatment. This data was benchmarked against *in vivo* transcriptome data from PF patients as indicated. **(b, c)** UMAP embedding of 63,581 single cells from the hPCLS model, color coded by cell type **(b)** and treatment **(c)**, respectively. **(d, e)** UMAP embedding of 481,788 single cells from the integrated multi-cohort PF cell atlas, color coded by cell type **(d)** and disease status **(e)**, respectively. **(f)** *Ex vivo* marker genes signatures of meta-cell types in hPCLS. The heatmap shows the average scaled expression of markers in each meta-cell type. **(g)** matchSCore comparison of *ex vivo* marker genes against *in vivo* marker genes. **(h)** Query-to-reference mapping with scArches overview: *ex vivo* hPCLS and *in vivo* PF scRNA-seq data were mapped to the Human Lung Cell Atlas (HLCA) reference for benchmarking and systematic side-by-side comparisons. **(i)** Similarity of CC and FC treated cells from hPCLS compared to cells from Control and Pulmonary Fibrosis in the PF-extended HLCA (*in vivo* reference) as assessed by scArches mapping. Stacked bar plots show the percentage of cells mapping to either cells from healthy controls or PF patients with regards to CC or FC treatment for each cell type. Fisher’s exact test. **(j)** IPF stage-specific signatures derived from publicly available bulkRNA-seq data^25^. The venn diagram illustrates the intersection of differentially upregulated genes in IPF stages 1, 2 and 3 (versus control tissue). Selected genes from these signatures and enriched GO terms are highlighted. **(k)** Comparison of hPCLS and IPF stage-specific signatures. The bar plot shows the overlap of differently upregulated genes (in %) between the *in vivo* stage-specific IPF signatures and the *ex vivo* hPCLS signature (FC vs. CC).

We hypothesized that treatment with FC would induce cell states that are also present in PF patients. To benchmark the hPCLS *ex vivo* model against *in vivo* transcriptomic changes observed in independent PF patient cohorts, we integrated our PF single cell data (Munich cohort^11^) with single cell data from three independent PF cohorts^8–10^ (Chicago, Nashville, and Newhaven) into an integrated PF cell atlas consisting of 481,788 single-cells from healthy and fibrotic lung tissues (143 patients, **Fig. 1d** and **e** and **Supp. Fig. 1h - m**). We annotated cell types in each dataset using a hierarchical approach and harmonized the annotations for comparison of the *in vivo* and *ex vivo* gene expression data. We defined 14 meta-cell types and 24 cell states in the *ex vivo* hPCLS data (**Fig. 1b** and **Supp Fig. 1a** and **c**) with distinct marker gene signatures (**Fig. 1f** and **Supp. Table S2-3**), which we found to be conserved in the *in vivo* reference (**Fig. 1g** and **Supp. Table S4-5**), irrespective of the experimental condition (**Supp. Fig 1n and o**).

Next, we leveraged query-to-reference mapping with scArches^26^ (**Fig. 1h**) to map both the *ex vivo* hPCLS data and the *in vivo* PF scRNA-seq data to the healthy integrated Human Lung Cell Atlas (HLCA)^27^ for a systematic side-by-side comparison of both data sources (**Supp. Fig. 2a and b**). We found that scArches maps FC-induced hPCLS cell states with a higher likelihood to PF patient-specific cell states than to controls (**Fig. 1i**, **Supp. Fig. 2c** and **d**), thereby validating that FC-treated hPCLS reproduce aspects of PF manifested in patients. The mapping showed that alveolar epithelial cells, capillary and vascular endothelial cells, as well as fibroblasts and smooth muscle cells/pericytes were more similar to cells from PF patients after treatment with FC compared to treatment with CC. (**Fig. 1i**). This was not the case for most immune cell types (except dendritic cells, Fisher’s exact test. **Fig. 1i**), hinting at possible limitations of the hPCLS model to fully recapitulate the immunological components of *in vivo* lung fibrogenesis.

**Figure 2.**
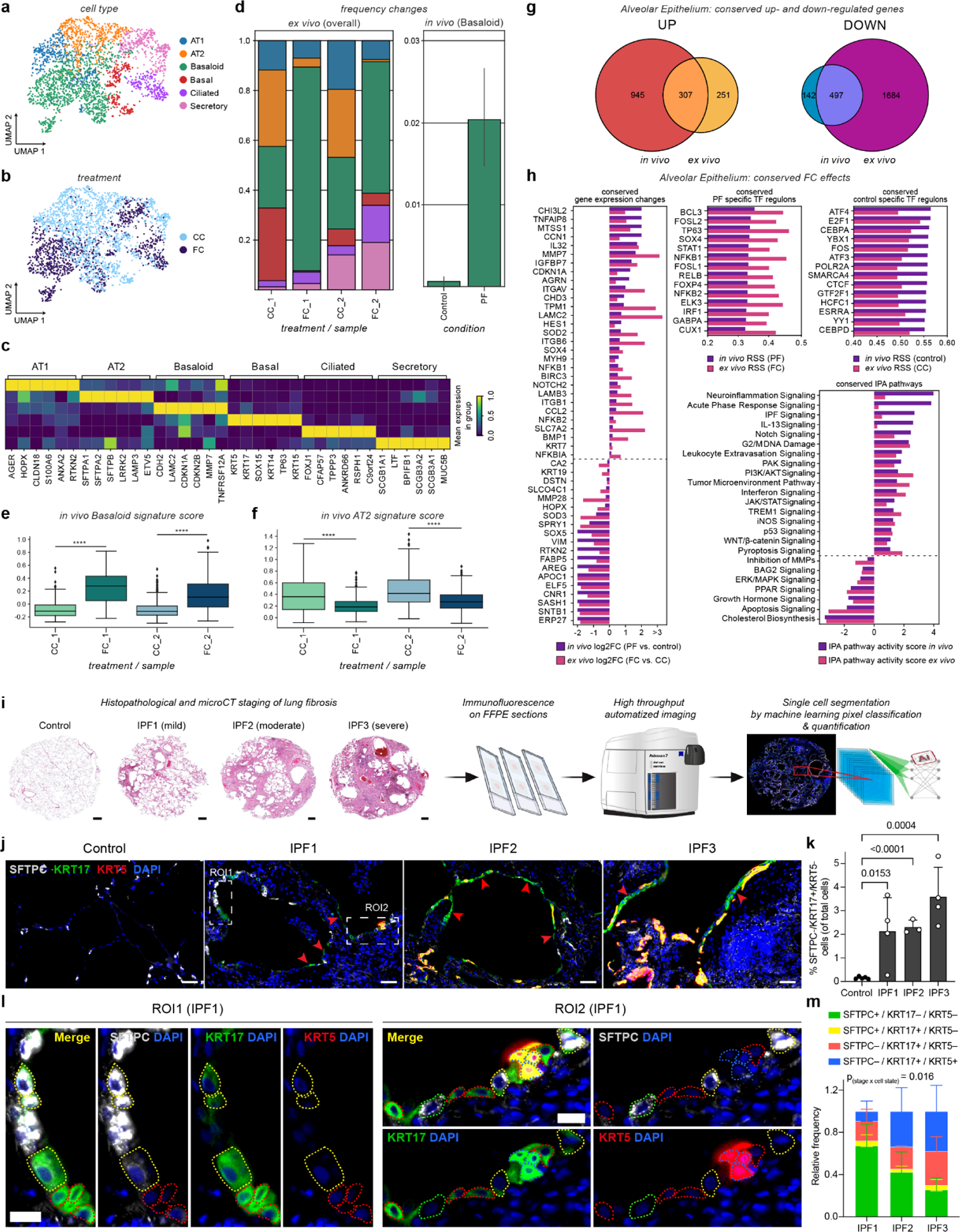
Induction of KRT17+/KRT5-basaloid cells in hPCLS *ex vivo* and early-stage IPF patient tissues. **(a,b)** UMAP embedding of 2,741 single epithelial cells from FC- and CC-treated hPCLS, color coded by cell type identity **(a)** and treatment **(b)**. **(c)** Ex vivo marker gene signatures of epithelial cells. The heatmap shows the scaled average expression in each cell type. **(d)** Cell type proportion analysis of epithelial cells in FC-vs CC-treated hPCLS (*ex vivo*, left) and PF vs control patients (*in vivo*, right): Stacked bar plot shows the relative frequency of each cell type with regards to treatment and tissue donor. **(e)** Boxplot of *in vivo* basaloid gene signature (derived from the *in vivo* reference PF cell atlas) score after treatment with FC or CC per tissue donor. Mann-Whitney-Wilcoxon test, ****p ≤ 0.0001. **(f)** Boxplot of *in vivo* AT2 cell gene signature (derived from the *in vivo* reference PF cell atlas) score after treatment with FC or CC per tissue donor. Mann-Whitney-Wilcoxon test, ****p ≤ 0.0001. **(g)** The Venn diagram illustrates the intersection of genes in alveolar epithelial cells that are uniformly upregulated both *in vivo* and *ex vivo*. **(h)** Qualitative analysis of conserved responses on the gene, pathway and Transcription Factor (TF) regulon level in alveolar epithelial cells. The bar plots illustrate the uniform behavior of conserved genes (log2FC), PF- and health-specific TF regulons (Regulon Specificity Score - RSS), and IPA pathways (IPA pathway activity score). **(i)** Staining and analysis pipeline assessing the presence of KRT17+/KRT5-basaloid cells by immunofluorescence (IF), high-throughput imaging and machine-learning segmentation of FFPE sections derived from histopathological and microCT staged lung tissues from IPF and control patients (scale bars = 1 mm). **(j)** Representative IF images of SFTPC (light gray), KRT17 (green) and KRT5 (red) stained lung sections demonstrating the presence of SFTPC-/KRT17+/KRT5-basaloid cells (red arrowheads) already in early-stage IPF (IPF1, scale bars = 50 μm). **(k)** Quantification of the percentage of SFTPC-/KRT17+/KRT5-basaloid cells by single-cell segmentations of 4 randomly selected regions of interest (ROIs) per patient (n = 5 controls, and n = 3 - 4 per IPF stage). In total, 249.700 single cells were analyzed. Statistics: Unpaired t-test. Bar plot depicts the Mean ± standard deviation. **(l)** ROI1 and ROI2 from IPF1 shown in panel **(j)** demonstrate distinct epithelial cell states: SFTPC+/KRT17-/KRT5-(green dashed contour), SFTPC+/KRT17+/KRT5-(yellow dashed contour), SFTPC-/KRT17+/KRT5-(red dashed contour) and SFTPC-/KRT17+/KRT5+ (blue dashed contour). Scale bars = 20 μm. **(m)** Stacked bar plots depicting the relative frequencies (Mean ± standard deviation) of the four distinct epithelial states across the different IPF stages. Statistics: Mixed effect model for two factors “stage” and “cell state”. p(stage x cell state) = 0.016. The same ROIs as in panel k and 25,466 single cells were quantified.

Transcriptomic states of pro-fibrotic macrophages have been described in recent single cell studies of PF patient lungs^8, 10, 12^. The scArches mapping did not identify significant changes in mapping for macrophages when assessing their overall distribution (**Fig. 1i**). Nevertheless, we identified several *in vivo/ex vivo* conserved aspects in macrophage states using more targeted analysis. Among 29,408 PTPRC+ immune cells in the hPCLS data, we identified 7,209 macrophages with marker gene sets that were conserved in the *in vivo* data (**Supp. Fig. S3 a-d** and **Supp. Fig. S4a-n**). While the overall frequency of macrophages was not altered after treatment with FC (**Supp. Fig. S3c**), we observed a distinct state of SPP1+ macrophages characterized by expression of fibrosis-associated marker genes such as SPP1, MMP9, CHI3L1, CCL2, and PLA2G7 (**Supp. Fig. S4d-j**), reminiscent of recently described macrophage states in PF and COVID-19^8–10, 12, 17^. Indeed, the *in vivo* SPP1+ macrophage signature was significantly increased in the *ex vivo SPP1*+ macrophage state (Mann-Whitney-Wilcoxon test, ****p ≤ 0.0001, **Supp. Fig. S4e**). Interestingly, the marker gene signature of this SPP1+ macrophage state appeared to share greater similarity with alveolar rather than interstitial macrophages (**Supp. Fig. S3f**). One-third of all genes upregulated and two-thirds of all genes downregulated in *in vivo* macrophages overlapped with *ex vivo* macrophage signatures at a statistically significant level (Wald test, adj. p-value < 0.05 and log2FC > 0.5; **Supp. Fig. S3g** and **Supp. Table S6**).

Of note, an important limitation of the hPCLS model is the lack of recruitment of circulatory monocytes that would give rise to fibrogenic macrophage states, possibly limiting the overlap of gene expression signatures. To functionally characterize the conserved gene expression shift in the *ex vivo* model, we performed IPA pathway analysis^28^, showing conserved induction of the Acute Phase Response, NFκ-B, IL1, and Cytokine Production by IL-17 pathways in the *ex vivo*model compared to PF patients (**Supp. Fig. S3h**). IL17, which can be induced by IL1, has been described to play a key role in causing pulmonary inflammation and fibrosis in both mice^29, 30^ and humans^31^. Interestingly, we also observed conserved induction of the IPA Pathway PD-1 / PD-L1 signaling (in cancer), which has previously been reported to play a role in the promotion of lung fibrosis^32, 33^. Transcription factor regulon activity analysis using SCENIC^34^ identified fibrosis-specific regulons, including STAT1, IRF1, IRF7, and CEBPB, that were opposed to the non-fibrotic regulons, which included KDM5A, IF8, and ZBTB7A (**Supp. Fig. S3h**).

We hypothesized that fibrogenesis in the FC-treated hPCLS rather reflects changes occurring early in disease progression. To analyze this aspect, we took advantage of a previously established micro-CT-based staging system that makes use of the spatial heterogeneity of end-stage IPF. Micro-CT staging allows the imaging-based stratification of lung samples according to their disease severity (mild = stage-1, moderate = stage-2, severe= stage-3) and thus serves as a proxy for analysis of disease progression without longitudinal sampling^25, 35^. We re-analysed bulk RNA-seq data from microCT staged tissue cores^25^ to compare the transcriptomic changes in the *ex vivo* model with changes at different stages of IPF progression *in vivo* (**Fig. 1a**). Four distinct gene signatures were extracted from the staged *in vivo* data; one specific to each of the three different stages of IPF progression and a ‘coré signature shared across all stages of IPF progression (**Fig. 1j**). Gene Set Enrichment Analysis (GSEA) on these four signatures revealed that the IPF ‘coré signature represented processes connected to cell adhesion and ECM deposition, including many well characterized marker genes of pro-fibrotic cell states (e.g. myofibroblasts and KRT17+/KRT5-basaloid cells). The IPF stage-1 signature was specifically enriched for myeloid immune processes and TGF-β1, the IPF stage-2 was enriched for lymphoid immune processes, and the IPF stage-3 signature for genes involved in cilium assembly and movement, likely reflecting an increase of ciliated cells, which have been shown to expand and line the bronchiolized epithelium of honeycomb cysts in IPF lungs^36, 37^.

We assessed the overlap of the four *in vivo* signatures with a ‘synthetic bulk’ signature derived from the *ex vivo* hPCLS single-cell experiments. The FC-induced gene expression changes in hPCLS were most similar to the *in vivo* core signature with 46% overlap, followed by an 11% overlap with the stage-1 specific *in vivo* signature, indicating that FC-treated hPCLS best recapitulates molecular processes that occur early in IPF progression and persist until end-stage disease (**Fig. 1k**).

In summary, our detailed single-cell map of *ex vivo* lung fibrogenesis in hPCLS highlights the power and limitations of the hPCLS organotypic model system. Our findings suggest that hPCLS treated with FC can be utilized to study the molecular and cellular mechanisms during the early phase of human lung fibrogenesis, which precedes the bronchiolization and honeycomb formation characteristic of end-stage IPF lungs.

### KRT17+/KRT5-basaloid cells appear in early-stage disease and in the *ex vivo* fibrosis model

Using longitudinal scRNA-seq analysis, we have previously shown that upon acute lung injury in mice both alveolar AT2 cells and airway-derived stem cells converge onto a highly distinct transitional cell state (Krt8+ Alveolar Differentiation Intermediate; Krt8+ ADI), which precedes their differentiation into AT1 cells during lung regeneration^18^. We and others have shown that this Krt8+ ADI cell state is characterized by gene programs that are reminiscent of epithelial to mesenchymal transition (EMT), cellular senescence and p53 activity, featuring expression of many well-known pro-fibrogenic paracrine secreted factors^18, 38–40^. Cross-species analysis of our mouse lung regeneration data with the PF cell atlas revealed that Krt8+ ADI cells were most similar to the KRT17+/KRT5-basaloid cells in PF patients^9, 10^, which also feature p53, EMT and senescence-like gene programs^39^.

We clustered 4,373 EPCAM+ epithelial cells in the hPCLS cytokine perturbation data (**Fig. 2a, b** and **Supp. Fig. S5a-c**) and identified alveolar AT1 and AT2 cells, as well as airway secretory, basal, and ciliated cell types based on canonical cell type markers (**Fig. 2c** and **Supp. Fig. S5d-s**). We also identified a large number of cells that were reminiscent of the KRT17+/KRT5-basaloid cells, which were specific to the diseased condition in the integrated PF cell atlas. The frequency of these KRT17+/KRT5-basaloid cells was significantly increased upon FC stimulation of the hPCLS (**Fig. 2d**). Using the *in vivo* marker signature of basaloid cells, we scored single cells in the hPCLS data and found a significant induction of the basaloid program in epithelial cells treated with FC compared to control conditions (**Fig. 2e**), which was accompanied by a significant reduction of the *in vivo* AT2 cell signature (**Fig. 2f**). Assuming a potential origin of basaloid cells in the alveolar epithelium we compared differential gene expression of alveolar epithelial cells *in vivo* and *ex vivo*, revealing a strong overlap in both up- and downregulated genes (55% of all genes upregulated *ex vivo* were also upregulated *in vivo*, while 23% of all genes downregulated *ex vivo* were also downregulated *in vivo*, Wald test, adj. p-value < 0.05 and log2FC > 0.5) (**Fig. 2g** and **Supp. Table S6**).

Among the conserved genes, we detected well-described markers of KRT17+/KRT5-basaloid cells such as MMP7, CDKN1A, ITGAV, and LAMC2, as well as the transcription factor SOX4, a regulator of EMT (**Fig. 2h**). Indeed, the regulon signature of SOX4 was among the top conserved TF regulons that were both highly specific for PF patients and FC-treated hPCLS (**Fig. 2h**), suggesting a key role of SOX4 during fibrotic remodeling of the alveolar epithelium. We further found that the regulon signature of ATF3, a regulator of mitochondrial homeostasis, was specific for the control condition both *in vivo* and *ex vivo* (**Fig. 2h**), which is in line with findings describing a protective and restorative role of ATF3 in the alveolar epithelium during lung injury in mice^41, 42^.

Ingenuity pathway analysis on epithelial changes confirmed ‘IPF signalinǵ as one of the top conserved pathways between *in vivo* and *ex vivo*, further validating FC-treated hPCLS as a model of fibrogenesis. The conserved gene signatures were further enriched for Notch signaling and WNT/β-Catenin signaling activity (**Fig. 2h**), which have both been shown to be aberrantly increased in AT2 cells in IPF, thereby impairing alveolar regeneration and promoting fibrosis^43–47^. We also noticed, however, that several pathways, including HGF and IL1 signaling showed opposite behavior *in vivo* and *ex vivo* (**Supp. Fig. 5t and u**), indicating that hPCLS might be an inadequate model to study their role in PF.

Having shown that the hPCLS model rather recapitulates early fibrogenic events, we next quantified the relative proportion of KRT17+/KRT5-basaloid cells amongst 249.700 single cells *in situ* using tissue sections from histopathological and microCT staged lung cores of control (n=5) and IPF patients (n=4) using immunofluorescence and AI-based image analysis (**Fig. 2i**). Consistent with our *ex vivo* findings, we found a significant increase of KRT17+/KRT5-basaloid cells already in IPF stage-1 in areas of mild alveolar septal thickening and AT2 cell hyperplasia, which exacerbated with advanced stages of IPF progression also propagating to histopathological regions of honeycombing and neo-bronchiolization (**Fig. 2 j,k**).

In conclusion, we demonstrate the appearance of KRT17+/KRT5-basaloid cells already in early-stage IPF and show that these cells can be induced in the *ex vivo* hPCLS model using pro-fibrotic cytokines.

### AT2 cells dedifferentiate towards a KRT17+/KRT5-basaloid cell state during *ex vivo* fibrogenesis

The cellular origin and function of KRT17+/KRT5-basaloid cells in PF is currently an ongoing debate, with some data suggesting a possible origin in AT2 cells^11, 48^. We indeed observed KRT5 negative cells that co-express the AT2 cell marker SFTPC and the basaloid cell marker KRT17 in areas of AT2 cell hyperplasia in the early-stage IPF tissues (**Fig. 2l, m**), suggesting that the AT2 cell may be a major source for generating basaloid cells in disease. To follow the fate of AT2 cells during *ex vivo* fibrogenesis, we quantified alveolar cell state transitions by immunofluorescence analysis on formalin-fixed and paraffin-embedded hPCLS sections treated with FC (**Fig. 3a**).

**Figure 3.**
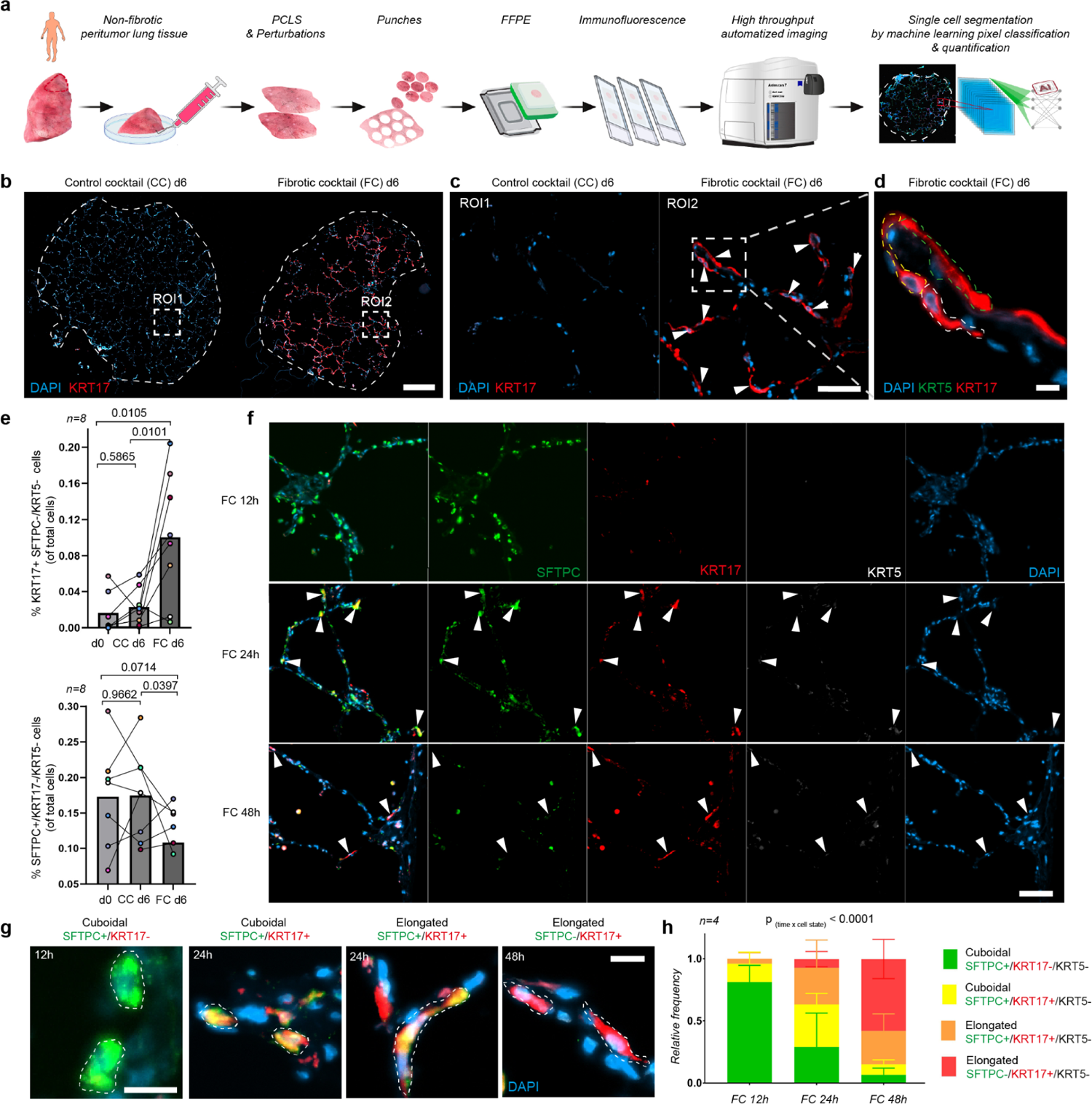
Time-resolved analysis shows AT2 cell conversion into KRT17+/KRT5-basaloid cells in hPCLS. **(a)** Staining and analysis pipeline using immunofluorescence (IF), high-throughput imaging and machine-learning segmentation of FFPE-punches taken from hPCLS. **(b)** IF analysis demonstrating an upregulation of KRT17+ expression (in red) at day 6 after FC compared to CC controls. Scale bar = 500 µm. **(c)** ROI1 and ROI2 from panel b represent a magnified view of CC-vs FC-treated hPCLS at day 6. Arrowheads within ROI2 indicate mostly elongated and KRT17+ cells (in red). Scale bar = 50 µm. **(d)** High-resolution view from panel c showing three single KRT17+ (in red) but KRT5-(in green) cells (circumvented in white, yellow, and green dashed lines). Scale bar = 15 µm. **(e)** Bar graphs show quantification by single-cell segmentations of entire PCLS FFPE-punches at day 6 after FC (n = 8 patients, n=130.630 single cells). Statistics: Unpaired t-test. **(f)** Time-resolved FC treatment (12h, 24h, 48h) of hPCLS and subsequent IF analysis staining for the indicated markers. Arrowheads within the FC 24h panel indicate KRT17+/KRT5-/SFTPC+ cells (red, gray, green) with morphology changes from cuboidal to elongated cells. Arrowheads within the FC 48h panel indicate elongated KRT17+/KRT5-(red/gray) cells with mostly negative SFTPC expression (green). Scale bar = 100 µm. **(g)** Four representative and distinct cell states derived from time-resolved (12h, 24h, 48h) FC treatments. Classification occurred according to morphology phenotypes (cuboidal or elongated) and KRT17/SFTPC expression after FC treatment over time. All of these cells were KRT5-. The KRT5-channel is not shown here. Scale bars = 15 µm. **(h)** Stacked bar plots depicting the relative frequencies (Mean ± standard deviation) of the four indicated cellular states measured at various time-points (12h, 24h, 48h) after FC treatment (n = 4 patients; n=1,136 single cells from 120 ROIs of 500 µm x 500 µm). Statistics: Two-way ANOVA calculated for two factors “time” and “cell state”. p(time x cell state) < 0.0001. All cell nuclei were stained with DAPI (blue).

On day 6 after FC treatment, we observed KRT17+/KRT5-cells reminiscent of basaloid cells with mostly elongated cell shapes covering the alveolar septa of hPCLS (**Fig. 3b-d**). Using single cell segmentation tools (Labkit and Imaris), we demonstrated a significantly higher proportion of KRT17+/KRT5-/SFTPC-cells in FC compared to CC, while the number of SFTPC+/KRT17-/KRT5-cells showed a significant reduction (**Fig. 3e** and **Fig. S6 a,b**). Importantly, these results validate our findings made by scRNA-seq (**Fig. 2**) in additional independent donor samples on the protein level (n=8 patients and n=130.630 single cells were analyzed).

The hPCLS model enables time-resolved analysis of the cell state shifts observed. We made use of this by fixing and sectioning CC- and FC-treated hPCLS punches at 12, 24, and 48 hours after treatment (**Fig. 3f-h**, n=4). At 12 hours after FC treatment, we mainly observed SFTPC+ *bona fide* AT2 cells with round cuboidal shapes. At 24 hours after treatment, a new state of AT2 cells emerged, which was still cuboidal but now co-expressed SFTPC and KRT17 while remaining KRT5 negative. Some of these SFTPC+/KRT17+/KRT5-cells showed more elongated cell shapes suggesting that the induction of the basaloid program may induce squamous differentiation in the AT2 cells. The number of these elongated SFTPC+/KRT17+/KRT5-cells was further increased at 48 hours after treatments, together with a reduction in the number of cuboidal SFTPC+/KRT17+/KRT5-cells. At 48 hours, we observed increased numbers of SFTPC-/KRT17+/KRT5-cells which also showed elongated squamous cell shapes (**Fig. 3f-h**, n=4).

Taken together, this data supports a model of AT2 cell differentiation towards KRT17+/KRT5-basaloid cells during the onset of human lung fibrogenesis. In contrast to *in vivo* observations that also featured alveolar KRT17+/KRT5+ basal cells already in stage-1 regions (**Fig. 2j and l**), we did not observe the generation of any basal cells after FC treatment in hPCLS.

### A basaloid-myofibroblast cell circuit in early-stage lung fibrosis is recapitulated *ex vivo* in hPCLS

KRT17+/KRT5-basaloid cells are potential paracrine regulators of fibroblast mediated fibrogenesis^9, 10, 18^. Indeed, our receptor-ligand analysis on the integrated PF cell atlas suggested a strong functional connection between basaloid cells and myofibroblasts (**Fig. S7a and b**). Myofibroblasts are central mediators of tissue fibrosis by secreting excess extracellular matrix (ECM). Single cell RNA-seq has recently helped to better characterize fibroblast heterogeneity in mouse and human lung fibrosis and identified CTHRC1+/TNC+ myofibroblasts as the major ECM secretion factory in lung fibrogenesis^19, 49^. To characterize fibroblasts and the myofibroblast state in hPCLS, we annotated the 20,761 COL1A2+ stromal cells contained in our data, leading to the identification of pericytes, smooth muscle cells (SMCs), fibroblasts, and myofibroblasts (**Fig. 4a, b**) with distinct marker genes conserved with their *in vivo* counterparts (**Fig. 4c** and **Fig. S8a-t**).

**Figure 4.**
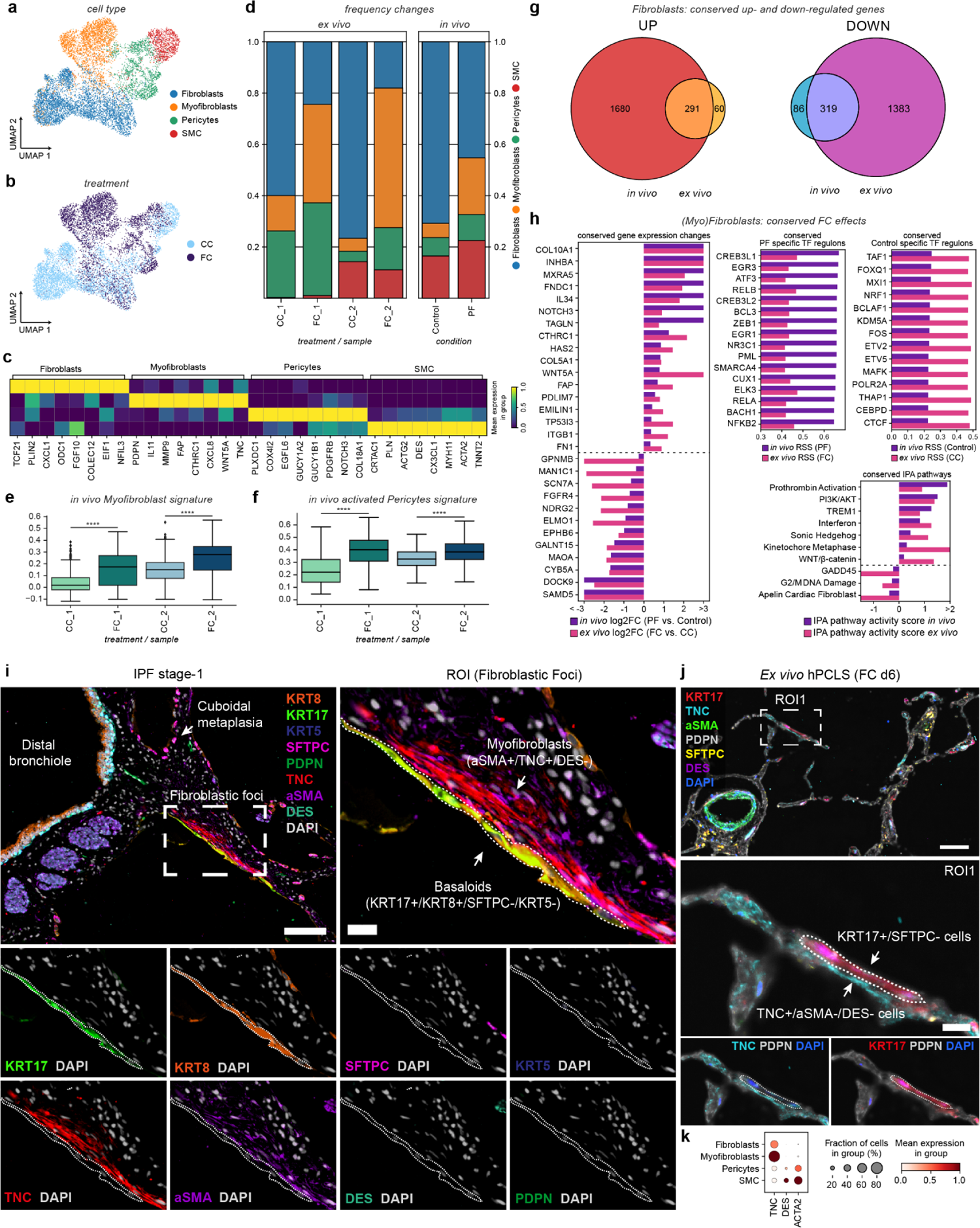
Myofibroblast-basaloid cell interactions in early-stage disease are recapitulated in the hPCLS model. **(a,b)** UMAP embedding of 9,721 single stromal cells from FC and CC treated hPCLS, color coded by cell type identity **(a)** and treatment **(b)**. **(c)** *Ex vivo* marker gene signatures of stromal cells. The heatmap shows the scaled average expression in each cell type. **(d)** Cell type proportion analysis of mesenchymal cells in FC-vs CC-treated hPCLS (*ex vivo*, left) and Pulmonary Fibrosis (PF) vs control patients (*in vivo*, right): Stacked bar plot shows the relative frequency of each cell type with regards to treatment and tissue donor and displays induction of myofibroblasts after treatment with FC. **(e)** Boxplot of *in vivo* myofibroblast gene signature score after treatment with FC and CC per tissue donor. Mann-Whitney-Wilcoxon test, ****p ≤ 0.0001. **(f)** Boxplot of *in vivo* activated pericyte gene signature (derived from Mayr et al., 2021^11^) score after treatment with FC and CC per tissue donor. Mann-Whitney-Wilcoxon test, ****p ≤ 0.0001. (g) Quantitative side-by-side comparison of genes differentially expressed in fibroblasts (fibroblast + myofibroblasts) *in vivo* and *ex vivo*. The Venn diagram illustrates the intersection of genes that are uniformly upregulated both *in vivo* and *ex vivo*. **(h)** Qualitative analysis of conserved responses to FC on the gene, pathway and Transcription Factor (TF) regulon level in fibroblasts (fibroblast + myofibroblasts). The bar plots illustrate the uniform behavior of conserved genes (log2FC), IPF- and health-specific TF regulons (Regulon Specificity Score - RSS), and IPA pathways (IPA pathway activity score). **(i)** Representative indirect iterative IF image (4i) of the indicated markers in fibroblastic foci of early-stage IPF (IPF1). Scale bars = 100 µm (overview image) and 20 µm (enlarged view). **(j)** The interaction of elongated SFTPC-/KRT17+ cells (white dashed line) and TNC+/aSMA-/DES-(myo-)fibroblasts is also recapitulated *ex vivo* in FC-treated hPCLS at d6. Scale bars = 100 µm (overview image) and 20 µm (enlarged view). (k) Dotplot showing the scaled average expression and fraction of stromal cells per group expressing *TNC*, *DES* and *ACTA2* (aSMA) in hPCLS.

**Figure 5.**
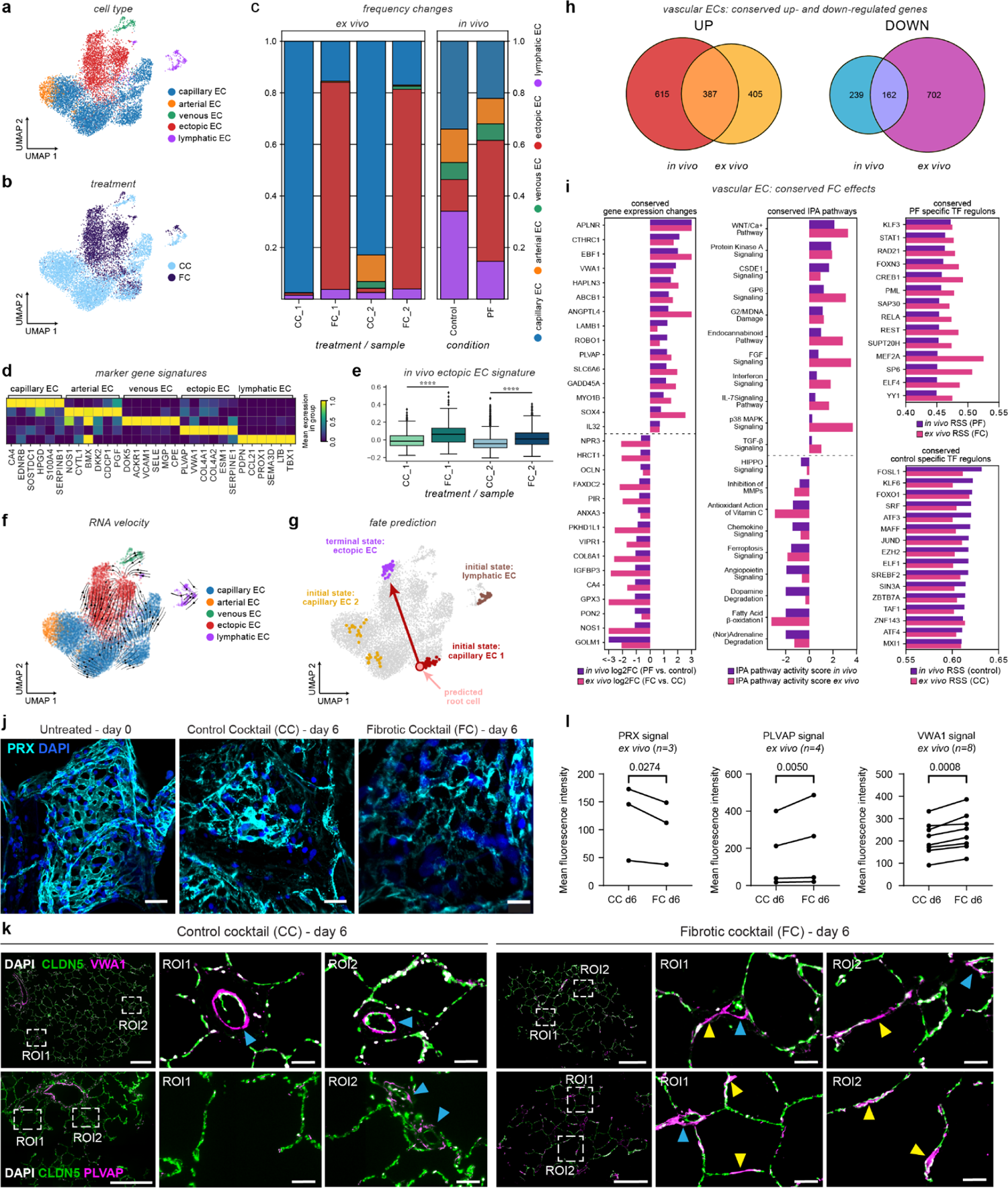
Emergence of a VWA1+/PLVAP+ EC state in hPCLS fibrogenesis. **(a,b)** UMAP embedding of 10,418 single endothelial cells (ECs), color coded by cell type identity and treatment. **(c)** Cell type proportion analysis of ECs reveals an expansion of ectopic ECs both in FC-treated hPCLS (*ex vivo*, left) and Pulmonary Fibrosis (PF) (*in vivo*, right): Stacked bar plot shows the relative frequency of each cell type with regards to treatment and tissue donor. **(d)** *Ex vivo* marker gene signatures of ECs. The heatmap shows the average scaled expression in each cell type. **(e)** Boxplot of the *in vivo* VWA1+/PLVAP+ EC cell gene signature score after treatment with FC with regard to treatment and tissue donor. Mann-Whitney-Wilcoxon test, ****p ≤ 0.0001. **(f)** RNA velocity projected on the UMAP embedding shows movement from capillary ECs towards ectopic ECs, indicating possible differentiation trajectories along the vectors. **(g)** CellRank^70^ fate mapping results projected on the UMAP embedding. CellRank identifies ectopic ECs as the terminal state of the differentiation process and infers three possible initial states, among which it predicts a subset of capillary ECs to be the most likely progenitor of ectopic ECs. **(h)** The Venn diagram illustrates the intersection of genes that are uniformly upregulated both i*n vivo* and *ex vivo*. **(i)** Qualitative analysis of conserved responses to FC on the gene, pathway and Transcription Factor (TF) regulon level across vascular ECs (arterial + venous + ectopic ECs). The bar plots illustrate the uniform behavior of conserved genes (log2FC), PF- and health-specific TF regulons (Regulon Specificity Score - RSS), and IPA pathways (IPA pathway activity score). **(j)** *Ex vivo* 3D whole mount immunofluorescence images of the indicated markers in hPCLS. Scale bar = 20 μm. **(k)** *Ex vivo* 2D immunofluorescence images of the indicated markers. Arrowheads in blue indicate pre-existing systemic venous ECs, while arrowheads in yellow indicate potential origin of ectopic vessels from capillaries. **(l)** Quantitative comparisons of mean alveolar fluorescence intensities of the indicated markers after 6 days of treatment. Statistics: Ratio paired t-test. Scale bars = 500 μm (overview images) and 50 μm (enlarged views).

Myofibroblasts in hPCLS were characterized by high expression of FAP, TNC, and FN1 in addition to the characteristic CTHRC1 marker gene (**Fig. 4c** and **Fig. S8d-l**) and were significantly increased in frequency upon treatment with FC (**Fig. 4d**). The *in vivo* myofibroblast signature was significantly increased after treatment with FC within COL1A2+ cells (Mann-Whitney-Wilcoxon test, ****p ≤ 0.0001, **Fig. 4e**). Similarly, we also found a significant induction of the activated pericyte signature we have previously described in lungs of IPF patients^11^ within COL1A2+ cells (Mann-Whitney-Wilcoxon test, ****p ≤ 0.0001, **Fig. 4f**).

The overlap of gene expression changes *in vivo* and *ex vivo* for fibroblasts was very high (83% for upregulated genes, Wald test, adj. p-value < 0.05 and log2FC > 0.5) (**Fig. 4g** and **Supp. Table S6**). Among the conserved genes, we found pathophysiologically relevant ECM genes COL10A1, COL5A1, EMILIN1, ITGB1, and FN1, indicating fibrotic ECM remodeling in treated hPCLS (**Fig. 4h**). We further identified NOTCH3, a mediator of myofibroblast differentiation^50^, and HAS2, which promotes invasiveness of myofibroblasts^51^, as key components of the conserved (myo)fibroblast gene program in hPCLS. Pathway enrichment analysis predicted increased activity of PI3K/AKT, Sonic Hedgehog (SHH), and WNT/ß-Catenin signaling pathways both *in vivo* and *ex vivo* (**Fig. 4h**), all of which were previously implicated in the pathogenesis of PF^52–56^. Our analysis also identified conserved Pro-Thrombin activation in the model, which has previously been shown to promote myofibroblast differentiation as well as collagen production^57, 58^. We further observed that HIF1-α signaling, which has been reported to be increased in fibroblasts from patients with PF and associated with increased tissue stiffness^59^, was among the pathways with discordant activity in hPCLS (**Supp. Fig. 8u and v**). As HIF1-α signaling was increased *in vivo* but showed decreased activity *ex vivo*, hPCLS appears to be an inept model for mechanistic studies of this particular pathway.

Through TF regulon analysis, we identified conserved TF regulons characteristic for the distinct fibroblast states in PF and control lungs. We observed fibrosis-specific induction of CREB3L1, which is a known upstream regulator of the transcription of COL1A1 and COL1A2^60^, as well as EGR1 and EGR3, both belonging to the Early Growth Response TF family, which has been shown to regulate TGF-β dependent fibrosis^61^ (**Fig. 4h**). TF regulon analysis using SCENIC further revealed ELK3 and NR3C1 as potential novel TF regulons of myofibroblast identity. In addition, we found the regulon signature of NRF1, a central regulator of cellular homeostasis during development^62, 63^, to be specific for the control condition both *in vivo* and *ex vivo*, suggesting a central role of NRF1 in maintaining the identity of homeostatic, non-pathological fibroblasts in the human lung.

To validate the predicted myofibroblast – basaloid crosstalk (**Fig. S7a and b**) *in situ*, we performed indirect iterative immunofluorescence imaging (4i)^64^ staining for eight different epithelial (KRT17, KRT5, KRT8, SFTPC, PDPN) and stromal (TNC, aSMA, DES) cell markers in stage-1 IPF tissue sections. This analysis revealed a polarization and direct physical interaction of TNC+ myofibroblasts with flattened KRT17+/KRT5-basaloid cells, seemingly forming nascent fibroblastic foci in the thickening alveolar wall of early-stage IPF (**Fig. 4i** and **Supp. Fig. S9a, b**). Notably, the physical interaction of TNC+ myofibroblasts and elongated KRT17+ basaloid cells in the alveolar septum was also recapitulated *ex vivo,* specifically in FC-treated hPCLS but not in the control condition (**Fig. 4j** and **Supp.Fig. S9c, d**). Interestingly, in the *ex vivo* hPCLS model, besides their interaction with TNC+/ACTA2-/DES-myofibroblasts, elongated SFTPC-/KRT17+/KRT5-basaloid cells were additionally found in the alveolar septum in direct proximity to TNC+/ACTA2+/DES+ cells (**Supp. Fig. S9c**), presumably representing myofibroblastic pericytes (**Fig. 4k**).

In summary, the spatial analysis demonstrates that the direct physical contact between basaloid epithelial cells and myofibroblasts in the thickening alveolar septum or early-stage PF can be modeled in hPCLS *ex vivo*.

### A capillary cell-derived VWA1+/PLVAP+ ectopic endothelial cell state during lung fibrogenesis

Alterations of the vasculature in PF are poorly studied and their contributions to disease pathogenesis remain elusive^10, 16, 65^. Recent single cell RNA-seq studies have identified a population of endothelial cells (ECs) in PF that express high levels of COL15A1, PLVAP and VWA1^10, 16, 65^. A similar set of marker genes has also been described to characterize the systemic venous circulation in healthy donor lungs^66^, raising the possibility that the systemic circulation expands in end-stage PF. Analysis of 16,513 ECs in the hPCLS data identified capillary ECs and clusters of arterial, venous and lymphatic ECs with correct expression of canonical markers (**Fig. 5a-d**).

Importantly, the FC perturbation induced a VWA1+/PLVAP+ ectopic EC state that was highly similar to the disease-associated cell state *in vivo* (**Fig. 5c** and **Supp. Fig. 10a-h and n-q**). Using the *in vivo* marker signature of VWA1+/PLVAP+ ECs, we scored all non-lymphatic endothelial cells in the hPCLS data and found a significant increase of this signature after treatment with FC compared to control conditions (**Fig. 5e**).

Interestingly and in contrast to the *in vivo* data, these cells were not expressing high levels of COL15A1. VWA1+/PLVAP+ ectopic EC also showed a uniquely high expression of COL4A1 and COL4A2 both *in vivo* and *ex vivo*, encoding subunits of collagen type IV, which is a core component of the basement membrane. Collagen type IV has been reported to be elevated in the serum of PF patients^67, 68^, suggesting ectopic ECs as a potential source of this serum biomarker.

The VWA1+/PLVAP+ ectopic ECs did not express marker genes of capillary (PRX, RGCC), arterial (BMX, VEGFA) or venous (VCAM1, ACKR1) ECs (**Fig. 5c**). Through comparative DGE testing, we recovered a set of features through which ectopic ECs distinguish themselves from normal vascular cells that are conserved in both PF patients and FC-treated hPCLS. 39% of all genes upregulated *in vivo*were also upregulated *ex vivo*, while 40% of all genes downregulated *in vivo* were also downregulated *ex vivo* (**Fig. 5h** and **Supp. Table S6**). Interesting examples were increased expression of ROBO1, a receptor for SLIT1 and SLIT2 that mediates cellular responses to molecular guidance cues in cellular migration, and the transcription factor SOX4 (**Fig. 5i**). VWA1+/PLVAP+ ectopic ECs also expressed higher levels of ANGPTL4, which has been shown to promote vascular leakage in the lung^69^, a hallmark of PF/acute lung injury. We further found conserved upregulation of WNT, PKA, FGF, IFN, IL7, and TGFß signaling pathways (**Fig. 5i**), suggesting that these pathways may play a central role in the emergence of VWA1+/PLVAP+ ectopic ECs in PF. However, our analysis also revealed the opposite regulation of BMP and JAK/STAT signaling (downregulated *in vivo*, upregulated *ex vivo*) (**Supp. Fig. 10r and s**), showcasing possible limitations of the hPCLS model.

We further found the TF regulon signature of KLF3, STAT1, RAD21, YY1 and CREB to be highly specific for both PF patients and FC-treated hPCLS (**Fig. 5i**). CREB has been shown to promote vascular integrity and has been linked to protective effects during acute lung injury in mice^71^, which may indicate a potential repair phenotype of VWA1+/PLVAP+ ectopic ECs. The ATF3 regulon signature was conserved in both control lungs and CC-treated hPCLS. Endothelial-specific loss of ATF3 has been shown to cause impaired alveolar regeneration^72^, suggesting a potential role in the alveolar homeostasis of the capillary endothelium.

Based on the loss of capillary ECs and simultaneous induction of the VWA1+/PLVAP+ state, we hypothesized that this EC state might be derived from capillary cells. We analyzed cell state trajectories using RNA velocity with scVelo^73^. Indeed, RNA velocity predicted a possible differentiation trajectory from capillary ECs towards VWA1+/PLVAP+ ectopic ECs (**Fig. 5e**). To make robust predictions of the lineage relationships that go beyond 2D projections of local RNA velocity vectors, we applied CellRank^70^, which allows for directional cell fate predictions by combining the directionality of RNA velocity vectors with trajectory inference algorithms. CellRank predicted VWA1+/PLVAP+ ectopic ECs as the terminal state of the FC-induced differentiation process and inferred three possible initial states. Analysis of the fate probabilities of all possible initial states showed that a subset of capillary ECs is the most likely progenitor of the VWA1+/PLVAP+ ectopic ECs (**Fig. 5f** and **Supp. Fig.10j-m**).

Next, we validated these findings using immunofluorescence analysis on both whole mount stainings and FFPE sections of the *ex vivo* treated hPCLS (n=6 donors), which revealed alterations in the capillary networks after FC treatment. FC treatment induced a significant loss of capillary PRX expression and integrity of the capillary network on day 6 after treatment (**Fig. 5j, l**). In control conditions, the expression of VWA1 and PLVAP was largely restricted to CLDN5+ vessels with apparent lumina, possibly representing systemic-venous endothelial cells described in the literature^66^. Importantly, in addition to these pre-existing VWA1+/PLVAP+ cells, we observed significantly increased de novo expression of VWA1 and PLVAP in the alveolar septa after FC treatment, further indicating that capillary ECs, that together with alveolar epithelial cells form the alveolar septum, dedifferentiate into the VWA1+/PLVAP+ ectopic EC state (**Fig. 5k, l**).

To analyze capillary EC plasticity *in vivo,* we used micro-CT staged IPF tissues (n=4), and quantified the emergence of VWA1+/PLVAP+ ECs in the alveolar septa of early-stage IPF (**Fig. 6**). PLVAP+ systemic vessels with large lumina were found in both control and IPF tissues across all stages. The number of PLVAP+ vessels showed a gradual increase with disease stage and increased luminal size compared to earlier stages (**Fig. 6a, b**). Correlating the thickness of the alveolar septum in healthy controls and early-stage IPF samples, we found significantly increased expression of PLVAP in the thickened alveolar septa of stage-1 IPF tissues compared to healthy control donors (**Fig. 6c, d**). Generally, the thicker the alveolar wall the more PLVAP expression was observed. A possible causal implication of VWA1+/PLVAP+ ectopic ECs with remodeling of the alveolar septum needs further investigation.

**Figure 6.**
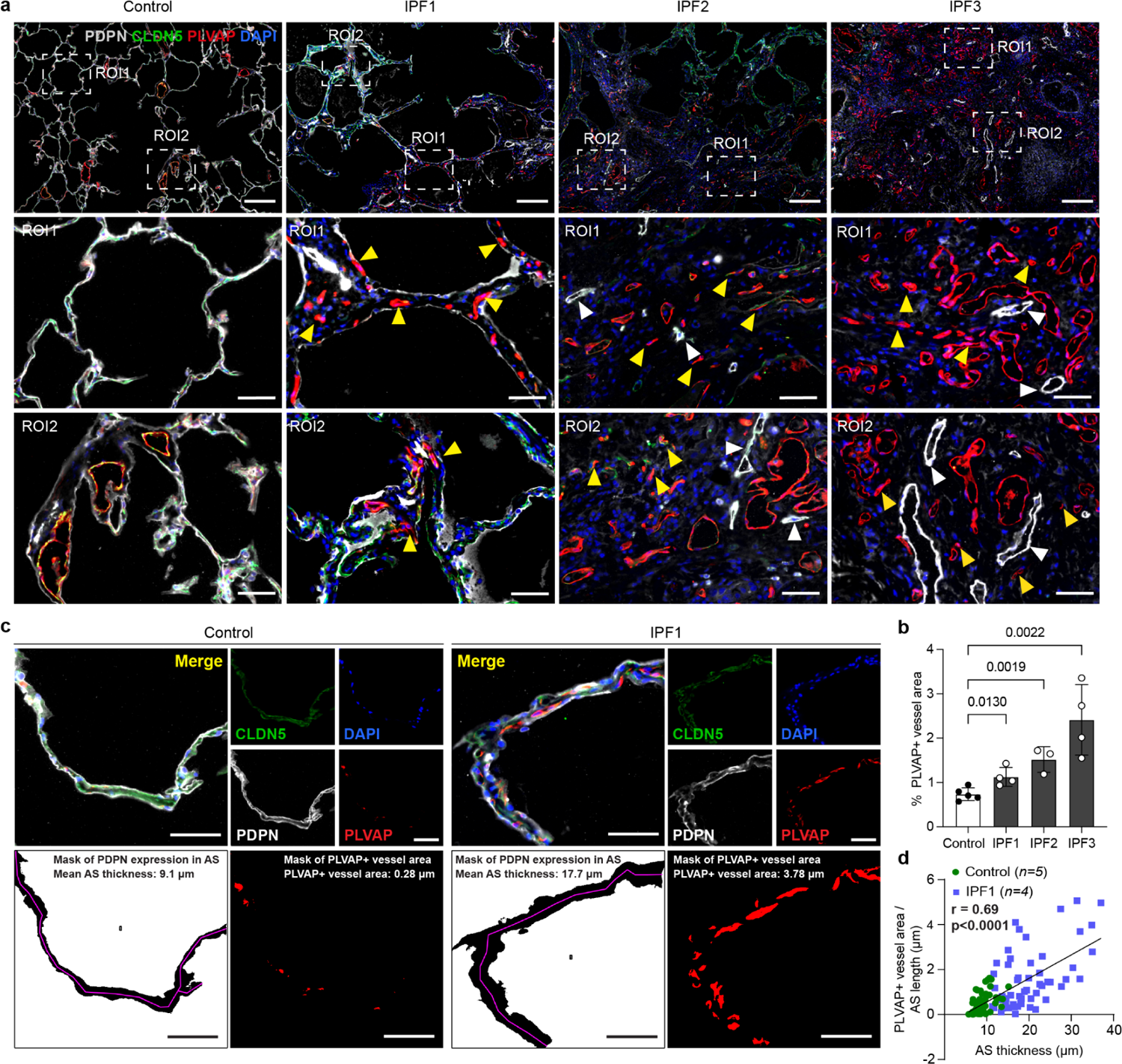
VWA1+/PLVAP+ ectopic ECs appear in alveolar septae of early-stage lung fibrosis. **(a)** Representative Immunofluorescence images of CLDN5 (green), PLVAP (red) and PDPN (white) expression in histopathological and microCT-staged IPF tissues. Yellow arrowheads indicate ectopic PLVAP+ vessels. White arrowheads mark PDPN+ lymphatics. Scale bars = 200 μm (overview images) and 50 μm (enlarged views). (**b)** Bar graphs show the quantification of PLVAP signal (percentage of PLVAP+ area to the total area of lung ROIs) *in vivo* (n = 5 control and n = 4 IPF patients). Statistics: Unpaired t-test. (**c)** Example images of quantification methodology of the alveolar PLVAP+ vessel area (normalized to the alveolar septal (AS) length) against the AS thickness. Representative alveolar septae of control and IPF1 tissues are shown. Scale bars = 50 μm. (**d)** The scatter plot shows the quantitative analysis of 60 AS derived from both control and IPF1 tissues, demonstrating a significant correlation of the normalized PLVAP+ vessel area with AS thickness (Pearson’s r =0.69, P<0.0001, n = 5 control and n = 4 IPF1 patients).

In summary, we have analyzed capillary EC plasticity during lung fibrogenesis *in vivo* and *ex vivo*. Thickened alveolar septa of early-stage IPF lesions showed loss of capillary cell identity together with a gain of the systemic circulation markers PLVAP and VWA1, revealing a possible capillary cell origin of the PLVAP+/VWA1+ ectopic ECs enriched in PF patients.

### A conserved fibrogenic alveolar cell-cell communication circuit in hPCLS

A unique strength of the hPCLS model compared to lung organoids is the presence of the full complexity of the lung’s cell types and 3D architecture that can be used to study patho-mechanistically relevant cell-cell communication. Therefore, we compared the cell-cell communication routes induced by FC in hPCLS to those observed in PF lungs. We used NicheNet^74^ to prioritize receptor-ligand pairs for each combination of sender and receiver cell type that are likely to regulate the perturbation effects and cell state changes in hPCLS and disease. Based on these predictions, we then calculated a directional interaction score which reflects how much each cell type contributed to the induction of the fibrogenic gene programs in the receiving cell types, taking into account the ligand activity score and the expression level of each ligand in one cell type relative to all other cell types (**Supp. Fig. S11a**). This approach allowed us to simultaneously dissect the directionality and quantify the strength of the cell-cell communication routes that drive fibrotic remodeling *ex vivo* and *in vivo*.

Besides vibrant crosstalk between structural cell types, we found myeloid cells (Macrophages and Mast cells, **Fig. 7a**) to be the most active immune cell types in the *ex vivo* network, whereas lymphoid cells (T and NK cells, **Fig. 7b**) played a more central role *in vivo*, indicating that the contributions of lymphoid cells to pathological lung remodeling during PF is not recapitulated in hPCLS. We speculate that the lymphoid cell crosstalk may occur only in later stages of PF or in the airway niche and hence is not captured in hPCLS by design, which would be in line with our findings that hPCLS is enriched for the IPF stage-1 signature. (**Fig. 1h**).

**Figure 7.**
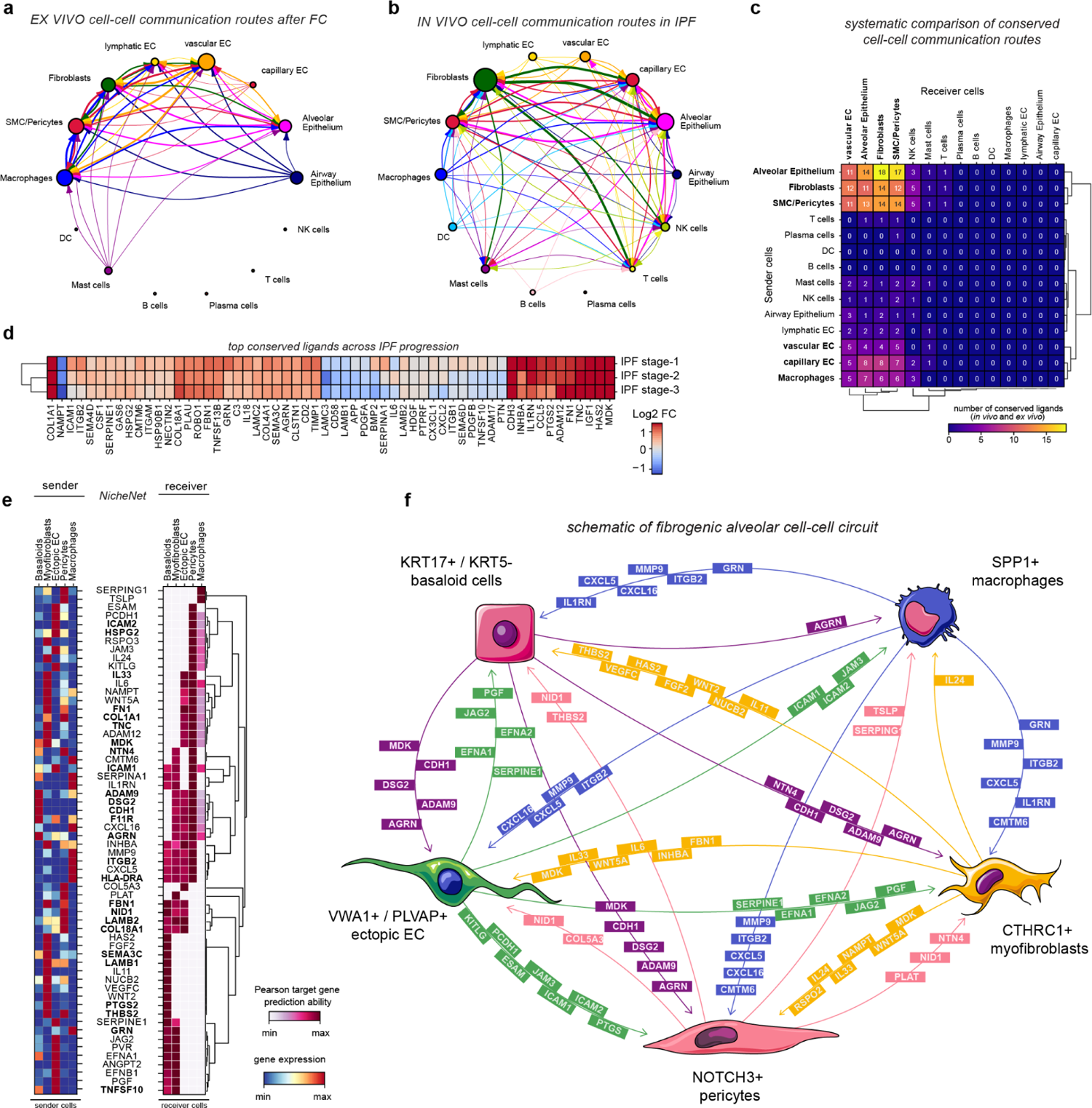
A conserved fibrogenic cell-cell communication circuit in hPCLS. **(a)** *Ex vivo* cell-cell communication routes in hPCLS after treatment with FC. Circle plot shows contributions of ligands upregulated after FC treatment (vs CC) in sender cell types to gene expression changes after treatment with FC (vs CC) in receiver cell types. The edge weight (thickness) represents the sum of the ligand scores, which is the product of the ligands expression in the sender cell and the NicheNet target gene prediction score. The color of the edge reflects the origin (sender cell) of the cell-cell communication route. The node size is proportional to the sum of the strength of outgoing cell-cell communication routes. **(b)** *In vivo* cell-cell communication routes in IPF. Circle plot shows contributions of ligands upregulated in IPF (vs healthy) in sender cell types to gene expression changes in IPF (vs healthy) in receiver cell types. **(c)** Number of conserved ligands (*in vivo* and *ex vivo*) from sender cells (rows) to receiver cells (columns) at meta-cell type level. The heatmap is clustered hierarchically and highlights a conserved cell-cell circuit between alveolar epithelial cells, vascular endothelial cells, fibroblasts, MCs/pericytes, and macrophages. **(d)** Variation of gene expression of the conserved ligands (*in vivo* and *ex vivo*) across IPF progression. Heatmap shows the log2FC of the identified conserved ligands IPF stages 1, 2 and 3 (vs healthy control tissue) as derived from publicly available bulkRNA-seq data (GEO GSE124685). **(e)** Overview of the cell-cell communication routes between five FC-induced cell states as predicted by NicheNet. The left panel shows the average expression of ligands predicted by NicheNet across sender cells. The right panel visualizes the corresponding predictive score of all ligands to induce their downstream target gene signature in the respective receiver cell types as measured by the Pearson correlation target gene prediction ability. Ligands in bold letters are conserved compared to *in vivo*. **(f)** Schematic of the pro-fibrotic cell-cell circuit. Ligands are derived from panel **(e)** and coloured according to the sender cell type they originate from.

Next, we aimed to quantify and identify which specific ligands between a pair of putative sender and receiver cells *in vivo* were conserved in the *ex vivo* model (**Fig. 7c** and **Supp. Fig. 11b and c**). Hierarchical clustering of the number of conserved ligands between identical sender and receiver cells *in vivo* and *ex vivo* revealed a conserved cell-cell circuit comprising alveolar epithelial cells, fibroblasts, pericytes/SMCs, endothelial cells and macrophages sustained by more than 60 conserved ligands, of which the vast majority had the highest bulk expression levels in IPF stage-1 and showed a gradual decline across IPF progression in stage-2 and 3 (**Fig. 7d**), indicating increased activity of this cell circuit in early-stages of IPF progression.

To better understand the interdependence of the disease-associated cell states that arise within this cell-cell circuit, we employed NicheNet^74^ to analyze the cell-cell communication routes between KRT17+/KRT5-basaloid cells, CTHRC1+ myofibroblasts, NOTCH3+ pericytes, VWA1+/PLVAP+ ectopic ECs, and SPP1+ macrophages (**Fig. 7f** and **g**). Out of the 60 ligands prioritized based on their capability to induce the specific transcriptomic signatures of the disease-associated cell states, 25 ligands were conserved between *in vivo* and *ex vivo* (**Fig. 7e** and **Supp. Fig. 11d**), conserved ligands highlighted in bold). Amongst the network of pairwise cellular interactions, we detected a conserved set of extracellular-matrix and cell-cell junction genes consisting of ADAM9, DSG2, CDH1, F11R and AGRN, specifically expressed by basaloid cells, that was characterized by its instructive role across the entire multi-lineage fibrogenic cell circuit.

To summarize, we delineated the cell-cell communication routes driving fibrotic remodeling in human lungs and hPCLS. A cell-cell communication circuit was largely conserved in hPCLS, enabling future disease relevant mechanistic perturbation studies.

### Drug-specific perturbation effects on the fibrogenic alveolar cell-cell communication circuit

Therapeutic options for the treatment of PF are limited and fail to halt disease progression. To establish the analysis of drug-specific perturbations on the pro-fibrotic cell-cell communication circuit in hPCLS, we employed a set of computational approaches that revealed drug-specific modes of action (**Fig. 8**). As a proof of concept, we analyzed the effects of the clinically approved anti-fibrotic drug Nintedanib^3^, as well as an *N*-(2-butoxyphenyl)-3-(phenyl)acrylamide (N23P) derivative of Tranilast (CMP4), which we recently have identified as a novel anti-fibrotic drug candidate^24^.

**Figure 8.**
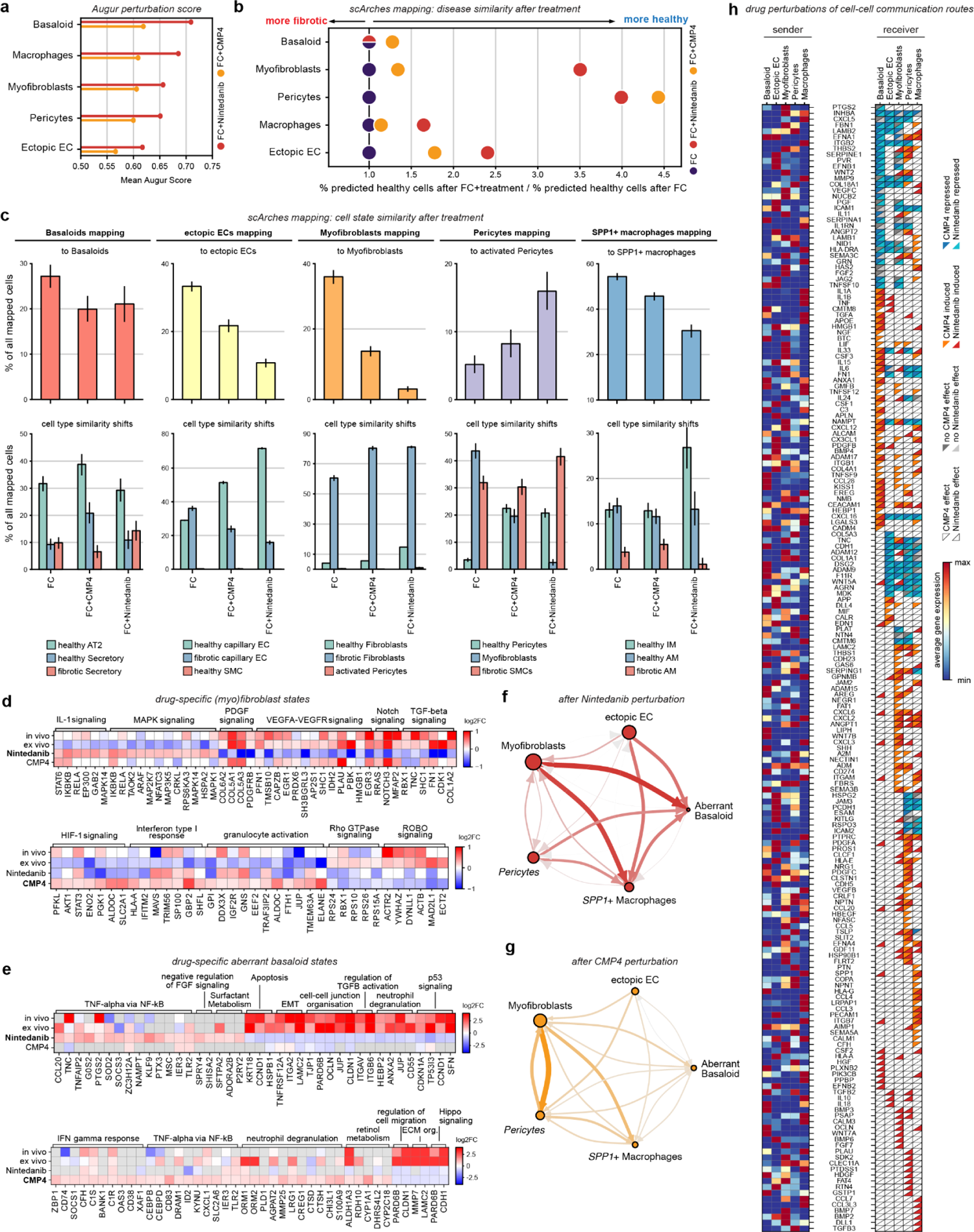
Cell lineage-specific effects and altered cell-cell communication upon drug treatments. **(a)** Bar graphs show the Augur perturbation score per cell state based on responsiveness to treatment with CMP4 or Nintedanib. **(b)** Drug effects on disease similarity as assessed through query-to-reference mapping of hPCLS data to PF-extended HLCA using scArches. The dot plot visualizes the mapping ratio after joint query-to-reference mapping of *ex vivo* hPCLS data to the PF-extended HLCA. **(c)** Drug effects on cell state identity shifts as assessed through joint query-to-reference mapping of *ex vivo* hPCLS data to the PF-extended HLCA. Bar plots show the percentage of cells mapped to each indicated cell state (top) and percentage of cells mapped to the top3 most similar cell types after the indicated treatments (bottom). Error bars show 95% confidence intervals. **(d,e)** Functional characterisation of drug-specific states of myofibroblasts **(d)** and basaloid cells **(e)**. Heatmaps show selected gene sets and genes identified through GSEA, which are specifically induced or repressed following treatment with either Nintedanib or CMP4. **(f,g)** Cell-cell communication routes after treatment with FC+Nintedanib **(f)** and FC+CMP4 **(g)**. Circle plot shows contributions of ligands upregulated after drug treatment (FC+CMP4/Nintedanib vs FC) in the respective sender cell state to gene expression changes in receiver cell types. The edge weight (thickness) scales with the cumulative strength of the ligands upregulated by the sender cell state to induce the observed gene expression changes in the sender cell type taking, into account the expression level as well as the NicheNet target gene prediction score of each ligand. The node size is proportional to the sum of all outgoing edges, reflecting the importance of the ligands to the cell-cell communication network upregulated in the respective cell state. **(h)** Perturbation map of drug-specific alterations of cell-cell communication routes. Left panel shows the average scaled expression of the ligands across sender cell states. Right panel illustrates whether the signaling route is unaffected, induced, or repressed by the treatment with the upper right triangle representing the Nintedanib effect and the lower left triangle representing the CMP4 effect. ECs: endothelial cells; AM: alveolar macrophages; IM: interstitial macrophages.

Drug treatments did not induce significant changes in cell type frequencies (**Supp Fig. 12a-e**). To quantify which cell states were most affected by the two treatments, we used Augur^75^ and found that Nintedanib introduces stronger changes in gene expression compared to CMP4 across all five fibrosis-specific cell states (**Fig. 8a**). Interestingly, KRT17+/KRT5-basaloid cells were most perturbed in response to both treatments, although both drugs are thought to primarily target myofibroblasts^24, 76^, possibly due to modulation of the cell-cell communication routes between myofibroblasts and basaloid cells by Nintedanib and CMP4.

We asked if the drug treatments shift cell states in a beneficial way towards a more ‘healthý-like transcriptional phenotype. To this end, we performed query-to-reference mapping using scArches^26^. We mapped hPCLS data to the integrated PF cell atlas and analyzed how drug treatments affected the proportion of nearest neighbor cells derived from healthy controls or PF patients in the reference (**Fig. 8b** and **Supp. Fig. 12f**). We found that following treatment with Nintedanib both myofibroblasts and pericytes mapped 3-4 times more often to cells from healthy controls donors in the reference, confirming its potential to revert fibroblast mediated fibrosis. CMP4 treatment had similar effects on pericytes, with a substantially lower effect on myofibroblasts. These drug effects were highly selective, with a slight improvement of healthy cell mapping for macrophages and ectopic ECs but very little beneficial effects on basaloid cells (**Fig. 8b**).

From these results, we concluded that a strong transcriptional perturbation of a cell state, in this case the basaloid cells (**Fig. 8a**), does not necessarily entail a substantial shift towards a more healthy cell identity (**Fig. 8b**). To achieve a more nuanced characterization of the treatment response that goes beyond the dichotomic distinction ‘healthy’ versus ‘fibrotic’, we additionally quantified shifts in cell type similarity induced by CMP4 and Nintedanib treatment using the same approach (**Fig. 8c**, **Supp. Fig. 12g-k**). We first analyzed whether the treatment led to relative changes in the similarity between the *ex vivo* cell states and their *in vivo* counterparts (**Fig. 8c**, top), followed by an unbiased overview of cell type similarity changes induced by the drug treatments (**Fig. 8c**, bottom). In contrast to Nintedanib, CMP4 treatment shifted the mapping of basaloid cells towards healthy AT2 cells in the HLCA reference by a factor of two (**Fig. 8c**).

We further observed significant reductions in the similarity of VWA1+/PLVAP+ ectopic ECs, CTHRC1+ myofibroblasts, and SPP1+ macrophages with their *in vivo* counterparts following CMP4 and Nintedanib treatment, indicating that both drugs dampen, but not reverse PF-associated cell state changes. Consistent with the Augur predictions, these effects were stronger after Nintedanib compared to CMP4 treatment. Both drug-induced transcriptional states of ectopic ECs exhibited a substantial increase in the similarity to healthy capillary ECs, albeit at different magnitudes, supporting our cell trajectory-based fate predictions (**Fig. 5**). While CMP4 induced no apparent cell type similarity shift of SPP1+ macrophages, the Nintedanib-induced cell state was substantially more similar to healthy interstitial macrophages (**Fig. 8c**).

Myofibroblast mapping to myofibroblasts in PF was substantially reduced by both drugs. The cells did, however, still map to cells from PF patients (‘fibrotic fibroblastś), indicating that even though the myofibroblast state got inhibited, a normal homeostatic fibroblast phenotype cannot be reached by both drugs. (**Fig. 8c**). A high fraction of the *ex vivo* NOTCH3+ pericytes mapped to myofibroblasts from PF patients, which was drastically reduced by Nintedanib treatment and also slightly reduced by CMP4. Interestingly, this went along with increased mapping to *in vivo ‘*activated pericyteś, a novel inflammatory pericyte state which we have previously shown to be associated with the degree of fibrotic remodeling (Ashcroft score) in PF patients^11^.

Next, we performed GSEA on all genes specifically induced or repressed by either of the two drugs and found, consistent with the literature, that Nintedanib inhibited VEGF and PDGF signaling in myofibroblasts^76^, providing proof of concept that the approach described here reveals correct mode of action data (**Fig. 8d** and **Supp. Table S6-7**). We further identified direct inhibition of TGF-β and Notch signaling as well as induction of IL-1 and MAPK signaling after Nintedanib treatment in fibroblasts.

The Nintedanib-induced changes in basaloid cells were characterized by suppression of some key features of these cells, including downregulation of several EMT genes (TNFRSF12A, ITGA2, LAMC2), TGF-β activation genes (ITGAV, ITGB6), and p53 signaling genes (TP53I3, CCND1, SFN) as well as a slight upregulation of genes involved in surfactant metabolism (**Fig. 8e**). These results appear to contradict our finding that Nintedanib does not significantly shift basaloid cells towards healthy cell identity (**Fig. 8b**). We think this demonstrates the potential of query-to-reference mapping, which harnesses the full gene expression space in an unbiased manner to classify the treatment response on a cellular level - in contrast to evaluating treatment response based on selections of established driver genes.

CMP4 exerted a pro-inflammatory effect on both myofibroblasts and basaloid cells mediated through induction of Interferon response and granulocyte activation/neutrophil degranulation in both cell states, as well as induction of TNF-α signaling genes in basaloid cells. Its inhibitory effects included less specific repression of Rho GTPase and Robo signaling in myofibroblasts and inhibition of ECM organization genes (MMP7, LAMC2) and Hippo signaling in basaloid cells (**Fig. 8d** and **e**).

Analyzing drug effects on cell-cell communication revealed drug-specific perturbation patterns in the conserved pro-fibrotic cell-cell communication circuit (**Fig. 8f-h** and **Supp. Fig. S12l**). Both drugs suppressed several predicted basaloid cell imprinting factors, including FBN1, THBS2, EFNB1, LAMB1, LAMB2, and NID1, while simultaneously inducing a new set of instructive ligands, dominated by interleukins IL1A, IL1B, IL33, IL15, IL6, and IL24, and TNF (**Fig. 8h**). Treatment with CMP4 led to the induction of several new signaling routes between myofibroblasts and pericytes; including NGF, LIF, and CCL5 from myofibroblasts and FAT1 from pericytes to myofibroblasts. On the other hand, Nintedanib treatment led to the emergence of BMP signaling routes via myofibroblast-derived BMP2 and BMP7 signals to SPP1+ Macrophages and basaloid-derived BMP3 signals to myofibroblasts and pericytes.

In summary, using query-to-reference mapping and cell-cell communication analysis, we identify cell type-specific drug modes of action and establish a roadmap for scalable drug testing in hPCLS.

## DISCUSSION

Modeling early-stage fibrotic disease directly in human tissue slice cultures offers transformative potential for both basic mechanistic studies as well as drug development and translation. In this work, we used single cell transcriptomics to carefully benchmark an *ex vivo* human precision-cut lung slice (hPCLS) model of lung fibrogenesis with *in vivo* patient data. Transcriptional changes and cell-cell communication routes in the *ex vivo* model closely reflected changes observed in the thickened alveolar septa of early-stage disease in idiopathic pulmonary fibrosis patients. Our analysis (1) reveals cell states and cell-cell communication routes in early-stage lung fibrogenesis as potential drug targets, (2) provides evidence for the cell type of origin for KRT17+/KRT5-basaloid cells and VWA1+/PLVAP+ ectopic ECs, (3) demonstrates cell lineage-specific drug effects within the alveolar cell circuit, and (4) serves as proof of concept for future drug mode of action and single cell perturbation studies in hPCLS.

Fibroproliferative forms of ARDS, as seen in COVID-19, feature very similar cell state transitions as we described here for PF^13, 16, 17^. For instance, our study highlights a so far underappreciated and poorly studied capillary EC plasticity upon human lung injury. We propose that pulmonary capillary EC in injured alveoli give rise to a VWA1+/PLVAP+ ectopic EC state with transcriptional similarities to the systemic vasculature. Increased numbers of these cells have been described previously in both PF^10^ and COVID-19^16^. Transcriptional similarities of the ectopic ECs in the lung to scar-associated endothelial cells in the human liver^77^ hint at a conserved mechanistic role shared across different entities of organ fibrosis.

The KRT17+/KRT5-basaloid cells are also prominently generated in both COVID-19 ARDS^13, 18^ and PF^9, 10^, suggesting that, similar to the myofibroblast and ectopic EC state, these basaloid cells can be conceptualized as Regenerative Intermediate Cell States (RICS) that normally resolve upon completion of lung regeneration. The cellular origin, future fate and pathophysiological relevance of basaloid cells in PF patients are currently unclear. Our time-resolved *ex vivo* perturbation data is consistent with an AT2-derived origin of basaloid cells, supporting recent work in lung organoids^48^. Our *in vivo* IPF stage-1 data features a heterogenous alveolar epithelium with hyperplastic AT2 and KRT17+/KRT5-basaloid cells that are occasionally interspersed with alveolar KRT17+/KRT5+ basal cells. It is possible that AT2 cell plasticity can generate these basal cells^48^; however, under the FC stimulation conditions described here, we did not observe the generation of KRT5+ cells in the alveolar space of hPCLS.

We can conceptualize the functional interactions of capillary EC, pericytes, alveolar epithelium and fibroblasts in the alveolar wall as an integrated cellular circuit that shifts its state during injury and fibrogenesis. The *in vivo* quantifications of micro-CT staged tissues show that basaloid cells and ectopic ECs appear in thickened alveolar septa of early-stage disease. In nascent fibroblastic foci, the myofibroblasts polarize towards basaloid cells and engage in direct physical contact. Importantly, we show that the functional cellular crosstalk of basaloid cells and myofibroblasts can be modeled *ex vivo* in hPCLS and present *ex vivo/in vivo* conserved candidate receptor-ligand pairs and transcriptional regulators of these cell state transitions. The anti-fibrotic drugs showed highly cell lineage-specific effects on these cell-cell communication routes, illustrating the importance of testing these drugs in tissue representing the full complexity of cell types.

Phenotypic drug discovery using single cell genomics is increasingly recognized by established industry and startup companies as an alternative paradigm to target-based approaches. Phenotypic drug discovery hinges on defining gene signatures (transcriptomic phenotypes) for initial (= lung fibrosis) and target (= healthy) tissue states that one is attempting to control. Phenotypic drug screens provide the basis for modeling the capability of screened drugs, or combinations thereof, to reach the target tissue state, reversing the initial disease state. Through a query-to-reference mapping approach, we demonstrate as a proof of concept that hPCLS coupled to perturbational single cell genomics can be used for phenotypic drug discovery by correctly identifying the known mode of action of the clinically approved drug Nintedanib. We anticipate that increasing the throughput of perturbational single cell genomics in this *ex vivo* model using emerging multiplexing techniques will enable accelerated drug development directly in human tissue and also form the basis for targeted analysis of gene regulatory networks, which we believe will be crucial for identifying drug candidates that exert high disease-reversing potency *in vivo*^78–80^.

We acknowledge several limitations of our work. Even though the main findings were extensively validated in a large number of additional donor samples using immunostainings, the scRNA-seq data was derived from two donor replicates. Given the high heterogeneity of human material used for PCLS, the future usage of novel scRNA-seq multiplexing methods will be required to enable higher replicate numbers cost effectively. Furthermore, as discussed, we view the hPCLS model used here as a model of fibrogenesis post-acute lung injury. This implies that likely not all genetic, environmental, and mechanistic factors that define a PF patient tissue are present, which may limit the power for specific drug effects that target aspects absent in the hPCLS model.

In conclusion, we demonstrate the induction of a multi-lineage circuit of fibrogenic cell states using pro-fibrotic cytokine stimulation in human lung tissue *ex vivo*. Computational query-to-reference benchmarking and *in situ* analysis of micro-CT staged patient tissues illustrates the relevance of this *ex vivo* model with respect to *in vivo* patient data. We anticipate that future experimental perturbations in hPCLS with single cell genomic readouts will be transformational for both basic science studies and the development and translation of drugs.

## Supporting information

Supplementary table S1: Baseline characteristics of included human IPF and control samples from Leuven

Supplementary table S2: Marker genes ex vivo (meta cell type level)

Supplementary table S3: Marker genes ex vivo (cell type level)

Supplementary table S4: Marker genes in vivo (meta cell type level)

Supplementary table S5: Marker genes in vivo (cell type level)

Supplementary table S6: Gene expression changes in vivo and ex vivo (in PF, after FC, FC+CMP4, and FC+Nintedanib treatment) for each meta cell type

Supplementary table S7: Gene expression changes of conserved cell-circuit cell states (basaloid, ectopic ECs, myofibroblasts, pericytes, SPP1+ Macroph

## Acknowledgements

We wish to thank all patients and their families who participated in this study. We gratefully acknowledge the provision of human biomaterial and clinical data from the CPC-M bioArchive and its partners at the Asklepios Biobank Gauting and the Klinikum der Universität München. We thank Anja Disovic and Marion Frankenberger from the CPC-M Bioarchive for excellent assistance in this project. We further thank Dr. Annette Feuchtinger and the Core Facility Pathology and Tissue Analytics at Helmholtz Munich for their excellent microscopy and imaging support, and Dr. Inti de la Rosa Velazquez and team at the Genomics Core Facility for sequencing and technical support. The authors are very grateful to Marisa Neumann and Anastasia van den Berg for their excellent technical assistance.

## Author contributions

HBS conceived and designed the study and supervised the entire work. GB and HBS supervised the hPCLS work. HBS, NJL, and JGS wrote the paper. DPG, MG and MW generated hPCLS and performed perturbation experiments. JGS, AA, CHM, and BHK generated scRNA-seq data. JGS, DPG, LY, RCJ, SZ, JCP, MW and YC performed and analyzed immunostainings. LJDS generated micro-CT staged IPF tissues. NJL generated custom code and led the analysis of scRNA-seq data. NJL, VS, MA, EG, VA and LH performed bioinformatic analysis of scRNA-seq data. FJT and MDL co-supervised the computational analysis. HBS, GB, FJT, MDL, MGS, WAW, JB, NK, RH, MG, and LS provided resources. All authors read and corrected the final manuscript.

## MATERIALS AND METHODS

### Human tissue and ethics statement

Fresh human tissue taken from tumor resections were obtained from the CPC-M bioArchive at the Comprehensive Pneumology Center (CPC Munich, Germany). The study was approved by the local ethics committee of the Ludwig-Maximilians University of Munich, Germany (Ethic vote #19-630). Written informed consent was obtained for all study participants.

Human lung IPF samples were provided from the KU Leuven lung biobank (ethical approval S52174). Samples were derived from explanted lungs, after written informed consent from all patients. Moreover, unused donor lungs were included as controls, following Belgian legislation. Four IPF lungs and five controls were included. Baseline characteristics of the included cases are provided in Supplementary table 1.

### Generation of human precision-cut lung slices (hPCLS) and treatments

Fresh tumor-free lung tissue was filled with a 3% (w/v) low melting agarose preparation media (Sigma, A9414-100G). Agarose filling was performed via a catheter inserted through one or several identified bronchi. Tissue blocks were cut in 500 µm thick PCLS using a vibration microtome Hyrax V50 (Zeiss) or a vibratome 7000smz-2 (Campden), as previously described^22^. hPCLS were cultured in DMEM-F12 (PAN Biotech, Cat. P30-3702) media containing 1% penicillin and streptomycin (Life technologies), and 1% amphotericin-B (life technologies) but deprived of serum. PCLS were incubated under 37°C, 5% CO_2_ and 95% humidity at all times. PCLS were treated in replicates of 2-4 per condition. For the day zero (d0) condition, the PCLS were freshly processed immediately after slicing, without further treatments. For all other time points PCLS were treated for 6 days with a control cocktail (CC) including all vehicles or a pro-fibrotic cocktail (FC) consisting of transforming growth factor-β (TGF-β) (5 ng/ml, Bio-Techne), platelet-derived growth factor-AB (PDGF-AB) (10 ng/ml, Thermo Fisher), tumor necrosis factor-α (TNF-α) (10 ng/ml, Bio-Techne), and lysophosphatidic acid (LPA) (5 µM, Cayman chemical) as described before^20, 23^. For the time-resolved analysis hPCLS were treated either with CC or FC, and hPCLS were collected either at d0, 12, 24, 48 hours, day 4, and day 6, washed in 1X PBS for sequencing or washed and fixed in 4% Paraformaldehyde (PFA) (Thermofisher) for 2 hours at 37°C for downstream processing for imaging. For drug treatments, hPCLS were treated with FC or CC, along with 10 µM Cmp4^24^ or 1 µM Nintedanib (Selleck, Houston, TX) for 6 days. All treatment solutions were replenished after four days with freshly prepared components.

### Generating single-cell suspensions from hPCLS

Two to four hPCLS slices at day 6 after CC, FC, FC+CMP4 and FC+Nintedanib treatment were washed in ice-cold 1X PBS and then cut into small pieces and enzymatically digested using a mixture of dispase (50 U/mL, Corning), collagenase (156 U/mL, Sigma), elastase (3.125 U/mL, Serva) and DNAse (17 U/mL, Qiagen) for 20 min at 37°C while shaking (750 rpm). Enzyme activity was stopped with 1X PBS supplemented with 10% FCS and dissociated cells were passed through a 30 µm cell strainer and pelleted by centrifugation (300 x g, 5 min, 4°C). When contamination with red blood cells was observed, red blood cell lysis with 1 ml 1x RBC lysis buffer (ebiosciences) for 1 min at room temperature was performed. After stopping the lysis with 1X PBS+10%FCS, cells were pelleted by centrifugation (300 x g, 5 min, 4°C) and subjected to dead cell removal (Miltenyi Biotec) according to the manufacturer’s instructions. After centrifugation of the live cell fraction (300 x g, 5 min, 4°C), cells were resuspended in 1X PBS supplemented with 1% FCS, and critically assessed for viability and counted using a Neubauer hemocytometer. For scRNA-seq, the cell concentration was adjusted to 1,000 cells/µl.

### Single-cell barcoding, library preparation, and sequencing

For scRNA-seq, around 16’000 cells (targeted cell recovery = 10’000 cells) were loaded on a 10x Genomics Chip G with Chromium Single Cell 3′ v3.1 gel beads and reagents (3′ GEX v3.1, 10x Genomics). Single index libraries were prepared according to the manufacturer’s protocol (10x Genomics, CG000204_RevD). After quality check, single-cell RNA-seq libraries were pooled and sequenced on a NovaSeq 6000 instrument (Read 1: 28 bp, i7: 8 bp, i5: 0 bp, Read 2: 91 bp).

### Preprocessing and analysis of *ex vivo* human PCLS scRNA-seq data

The generation of count matrices was performed using the Cellranger computational pipeline (v3.1.0, STAR v2.5.3a). The reads were aligned to the hg38 human reference genome (GRCh38.99). After preprocessing, analysis of the *ex vivo* human PCLS scRNA-seq data was conducted using the Python package Scanpy^81^ (version 1.8.2). To minimize the possible effect of potential batch correction methods on our annotation, we first processed and annotated the data separately for both biological replicates, before processing and annotating them jointly. During QC we removed all genes with fewer than 1 count and those that were expressed in less than 5 cells. Further, we removed all cells with greater or equal than 10% mitochondrial counts, fewer than 300 genes, or fewer than 500 total counts. Next, we performed doublet detection with Scrublet^82^ on the individual sample level, ambient RNA correction with soupX^83^ (background contamination fraction was manually set to 0.3), and scran’s size factor based normalization^84^ using cell groupings obtained through Louvain cluster (resolution = 2.0) as input. We then performed log transformation via Scanpy’s pp.log1p() function. Highly-variable genes (HVGs) were computed for each sample separately (top 4000 genes per sample) and only considered as overall HVGs, if they were highly variable in at least two samples. We obtained 1856 HVGs after removal of cell cycle genes. We then performed Principal Component Analysis (PCA) using the HVGs as input and constructed a kNN graph using the first 50 principal components. We then performed graph-based clustering using the Leiden algorithm^85^ and annotated the resulting clusters based on canonical marker genes. In the first step, we annotated the data coarsely and divided it into an EPCAM+ subset of epithelial cells, a COL1A2+ subset of stromal cells, a CLDN5+ subset of endothelial cells, and a PTPRC+ subset of immune cells. Subsequently, we repeated HVG selection, PCA, and graph based clustering to achieve a fine grained annotation of the four lineage subsets. Marker genes were computed using a Wilcoxon rank-sum test, and genes were considered marker genes if the FDR-corrected p-value was below 0.05 and the log2 fold change was above 0.5.

### Generation of the *in vivo* reference: multi-cohort integrated Pulmonary Fibrosis cell atlas

No new *in vivo* scRNA-seq data of human patient lungs were generated for this manuscript. We generated a multi-cohort integrated cell atlas of pulmonary fibrosis comprising four different studies which explored the transcriptomes of non-diseased and fibrotic human lungs^8–11^. To assess and address potential batch effects, for each cohort the publicly available raw count matrices were re-processed separately in a similar manner using the python package Scanpy^81^ (version 1.8.2). To mitigate effects of background mRNA contamination, we performed ambient mRNA correction using the adjustCounts() function from the R library SoupX^83^ (background contamination fraction was manually set to 0.3). We then normalized the resulting expression matrices with scran’s size factor based approach^84^ using cell groupings obtained through louvain clustering (resolution = 2.0) as input. Finally, we performed log transformation via Scanpy’s pp.log1p() and cohort-wise scaling to unit variance and zero mean of the separate count matrices before concatenating the separate count matrices into one final count matrix encompassing all four cohorts.

To identify a shared set of highly variable genes (HVGs) was selected we calculated gene variability separately for each patient/sample (top 4000 genes per sample). A gene was considered as variable if it was listed as highly variable across a minimum number of patients in the respective cohort. The thresholds were motivated by sample size of each cohort (Munich 4 patients, Chicago 5 patients, Nashville 8 patients, Nashville 10 patients). Finally, the intersection of the HVGs sets yielded 1311 final genes after removal of cell cycle genes. These genes then provided the input for the principal component analysis (PCA). A batch-corrected neighborhood graph, accounting for the different cohorts in the visualization, was calculated using the bbknn package^86^, defining the cohorts as batch key (n_neighbors = 20, n_componentsn = 50). For cell type annotation, we performed graph-based clustering using the Leiden algorithm to split the data and into an EPCAM+ subset of epithelial cells, a COL1A2+ subset of stromal cells, a CLDN5+ subset of endothelial cells, and a PTPRC+ subset of immune cells, which we then clustered recursively and annotated based on canonical marker genes from the literature. Analog to the *ex vivo* data, marker genes were computed using a Wilcoxon rank-sum test, and genes were considered marker genes if the FDR-corrected p-value was below 0.05 and the log2 fold change was above 0.5

### scArches mapping to benchmark *ex vivo* PCLS scRNA-seq data

We utilized scArches^26^ (version 0.5.5) to map our *ex vivo* hPCLS scRNA-seq data as well as publicly available *in vivo* PF scRNA-seq data to the integrated Human Lung Cell Atlas^27^ (HLCA) for benchmarking and systematic comparisons. We then mapped publicly available *in vivo* PF sc-RNAseq data to the HLCA which included various PF entities (annotated as “IPF”, “Systemic sclerosis-associated ILD”, “End-stage lung fibrosis, unknown etiology”, “Myositis-associated ILD” or “ILD” in the HLCA metadata). The atlas contained 922,332 cells in total, with 584,944 cells from healthy lung tissue and 337,388 cells from lungs with pulmonary fibrosis. The mapping of *ex vivo* hPCLS data was performed using 500 surgery epochs with Kullback-Leibler divergence weight equal to 0.5. Other parameters were set as recommended by the developers of scvi-tools^87^. Cell types and condition labels were transferred from the HLCA using the K-nearest neighbors classifier with K equal to 50. We used Fisher’s exact test with Bonferroni correction as implemented in R to evaluate statistically significant treatment effects on transferred cell type (**Supp. Fig. 11b - f**) and condition similarities (**Supp. Fig. 11a**). We define cell type similarity as the proportion of each manually annotated cell type in *ex vivo* hPCLS data that has been classified as a specific cell type by label transfer from the HLCA and analog for condition similarity. To evaluate the effect of CMP4 and Nintedanib treatment on, we calculated the ratio of the proportion of cells that were labeled as healthy by label transfer from the HLCA after FC+treatment to the proportion of cells that were labeled as healthy by label transfer from the HLCA in the FC group (**Fig. 8b**). Confidence intervals were obtained by bootstrap with 1000 iterations. To understand the proximity of cell types found in *ex vivo* hPCLS data to cell types from HLCA, we calculated centroids in latent embedding space for each cell type and sample, and visualized them on 2D PCA plot (**Supp. Fig. 2**).

### Re-analysis of publicly available bulkRNA-seq data from staged IPF tissue

To compare the *ex vivo* PCLS scRNA-seq data against publicly available *in vivo* bulkRNA-seq from IPF stages 1, 2, and 3 (GEO GSE124685) we generated ‘synthetic bulk’ data from our single-cell data. Therefore, we summed up the counts of each gene across all cells from each sample (prior to any quality control steps). Next, we performed differential gene expression testing with DEseq2^88^ (version 1.34.0) for both the *in vivo* bulkRNA-seq (IPF stage 1 vs. Control, IPF stage 2 vs. Control, IPF stage 3 vs. Control) and for the *ex vivo* ‘synthetic bulk’ data (Fibrotic Cocktail vs. Control Cocktail). Prior to testing, we excluded genes that had fewer than 10 counts across all stages. We considered genes differentially upregulated at a statistically significant level if the FDR-corrected p-value was below 0.05 and the log2 fold change was above 0.25.

### Differential gene expression testing (DGE)

Differential gene expression (DGE) testing was conducted on SCRAN normalized counts using diffxpy^89^ (version 0.7.4). We then performed a Wald test with default parameters to compare gene expression in two different groups for all genes that were expressed in at least 5 cells. We considered genes differentially expressed at a statistically significant level if the FDR-corrected p-value was below 0.05 and the log2 fold change was above 0.25 or below - 0.25.

### Gene signature scoring

Gene signatures were derived from the integrated PF cell atlas. The gene signatures comprise all marker genes of the respective cell type or state that fulfilled the following criteria: FDR-corrected p-value below 0.05, log2 fold change above 2, expressed by fewer than 25% of the background. The respective scores were then calculated by subtracting the average expression of randomly sampled genes from the average expression of the signature using Scanpy’s *sc.tl.score_genes()* function with default parameters.

### Gene Set Enrichment Analysis (GSEA)

Gene Set Enrichment Analysis was conducted using GSEApy^90^ (version 1.0.0) with default parameters using the the following libraries from the human database: ’MSigDB_Hallmark_2020’, ’GO_Biological_Process_2021’, ’GO_Cellular_Component_2021’, ’GO_Molecular_Function_2021’, ’Reactome_2016’, ’KEGG_2021_Human’. Gene sets were considered enriched in the respective signature at a statistically significant level if the FDR-corrected p-value was below 0.05.

### Ingenuity pathway analysis (IPA)

Conserved and perturbed pathways and upstream regulators were inferred through the use of IPA^28^ (QIAGEN Inc., https://www.qiagenbioinformatics.com/products/ingenuitypathway-analysis) using the results of the conserved or treatment-perturbed genes obtained by DGE testing (see above) as input.

### Transcription factor regulon activity analysis with SCENIC

Conserved transcription factor (TF) regulons (i.e. Pulmonary Fibrosis lungs / FC treated hPCLS) were identified using the R package SCENIC^34^ (version 1.1.2) with default settings. The top conserved TF regulons were identified based on the Regulon Specificity Score (RSS).

### Treatment effect analysis

To dissect the treatment effects of Fibroblasts, we first performed DGE testing in different settings (Pulmonary Fibrosis vs. Control, FC vs. CC, FC+CMP4 vs. CC, FC+Nintedanib) as described above. We then identified those genes among all genes upregulated (downregulated) both *in vivo* (IPF vs. healthy) and *ex vivo* (FC vs. CC), that were specifically repressed (induced) after treatment with either of the two drugs. Next, we performed GSEA as described above to functionally characterize the treatment induced or suppressed changes in gene expression.

To identify cell states most perturbed by drug treatments, we employed the Augur algorithm^75^ as implemented in the Python package pertpy^91^ (version 0.2.0) on the five fibrosis-associated cell states (parameters: n_subsamples=100, subsample_size=20, select_variance_features=True). Augur utilizes a classifier to assess how accurately the experimental perturbation labels (FC and FC+Nintedanib/CMP4) can be predicted for a single cell based on its gene expression pattern. The predicted perturbation label of a cell is then compared with its experimental perturbation label, which enables cell type prioritization based on the area under the receiver operating characteristic curve (AUC).

### Cell-cell communication analysis

Cell-cell communication analysis was performed using the R package NicheNet^74^ (version 1.0.0). A schematic of the workflow is outlined in **Supp. Fig S10a**. NicheNet requires users to define a set of sender cell types, one single receiver cell type and a gene set of interest, that is potentially induced in the receiver cell type through niche signals from the sender cell types. Therefore, we performed differential gene expression (DGE) testing (see above) for each potential receiver cell type to define perturbed gene programs after Fibrotic Cocktail (or any of the two drug treatments). Genes were included in the perturbed gene programs, if the FDR-corrected p-value was below 0.05, the log2 fold change was above 0.25, and the gene was expressed by at least 10% of receiver cells in the perturbed condition. For each receiver cell type the respective perturbed gene programs were then used as input for NicheNet to predict ligands that could induce the gene program in the receiver cell type. We only considered ligands, that were expressed by at 10% of sender cells, and had a pearson correlation prediction ability (with regards to the perturbed gene programs) was greater than 0.05. To measure the individual contribution of the predicted ligands expressed by each sender cell type to the induction of the perturbed gene program in the receiver cell types, we then calculated for each sender-receiver cell type pair the sum of the products of the pearson correlation prediction ability of each ligand and the average expression of the ligand in the respective sender cell type. These so-called directional interaction scores in their entirety allowed the generation of a cell type-cell type interaction matrix, which was used to quantify and visualize the directionality of cell-cell communication routes after perturbation.

### Trajectory inference analysis and fate mapping

Trajectory inference analysis and fate mapping was conducted using the Python packages scVelo^73^ (version 0.2.4) and CellRank^70^ (version 1.5.1). First, we inferred the splicing kinetics of all genes and estimated RNA velocity using the dynamical model in scVelo. Second, we computed macrostates, set injury induced cell states (such as ectopic ECs) as terminal states, and computed initial states using CellRank. Finally, we computed absorption probabilities per cell in order to calculate average fate probabilities for each initial state.

### Immunofluorescence (IF) stainings of PCLS, microscopy, and quantification

Human PCLS collected from a time series of CC or FC treatments (12h, 24h, 48h, 4d, and 6d) and untreated samples were stained in whole-mount or in 6-8 µm thick formalin-fixed paraffin-embedded (FFPE) tissue sections. Whole-mount immunostaining for PRX was performed as previously described^92^: hPCLS samples were blocked and permeabilized in PBSGT (1X PBS with 0.2% gelatin, 0.5% Triton X-100, 0.01% thimerosal) for 3-4 h at room temperature (RT). Next, the samples were incubated with a primary PRX antibody (Sigma, HPA001868, 1:500) in PBSGT + 0.1% saponin (10 µg/mL) overnight at 4°C and rinsed in PBS 1X + 0.5% Triton (PBST), 4 times during 20 mins with rotation. The secondary antibody and DAPI in PBSGT + 0.1% saponin (10 µg/mL) were incubated for 2-4 h at RT and followed by 3 washes in PBST at RT with rotation.

FFPE sections of punched hPCLS were first dried at 60°C for 1 hour, followed by tissue deparaffinization using two times Xylene for 3-5 min. Next, FFPE sections were rehydrated with a series of ethanol solutions (2x 100%, 90%, 80%, 70%) for 2-5 mins each and washed in 1X PBS. Afterwards, antigen retrieval with 10 mM citrate buffer, pH=6.0 in a pressure cooker (30 s at 125 °C and 10 s at 90 °C) was performed. 10% normal donkey serum or 1% BSA were used for blocking nonspecific binding and primary antibodies were incubated (VWA1 1:100, Proteintech, 14322-1-AP; CLDN5 1:100, Santa Cruz, sc-374221; PLVAP 1:300, NovusBIO, NBP1-83911; KRT5 1:2000, biolegend, 905901; KRT17 1:400, Sigma, HPA000453; SFTPC 1:50, Santa Cruz, sc-518029) overnight at 4°C. Secondary antibodies (donkey anti-mouse Alexa Fluor 488, A21202, 1:250; donkey anti-rabbit Alexa Fluor 647, A31573, 1:250, goat anti-chicken Alexa Fluor 568, A11041, 1:250; Goat anti-chicken Alexa Fluor 488, A11039, 1:500; donkey anti-rabbit Alexa Fluor 568, A10042, 1:500, goat anti-mouse Alexa Fluor 647, A21236, 1:500, all from Invitrogen) and DAPI were then incubated for 2 hours at RT.

Whole-mount stained hPCLS and FFPE sections were imaged using a confocal laser scanning microscopy (LSM 710, Zeiss) or an upright fluorescence microscope (AxioImager, Zeiss), equipped with an Axiocam and by using a Plan-Apochromat 20×/0.8 M27 objective. The automated 3D-imaging of total FFPE-punches from multiple slides was accomplished using an Axioscan 7 (Zeiss) and a Plan-Apochromat 20×/0.8 M27 objective. The final images were analyzed, stitched, cropped and individually adjusted for contrast and brightness by using the ZEN2012 (Zeiss) software.

Quantification of the expression of endothelial markers PRX, PLVAP, and VWA1 was performed in ImageJ (https://imagej.nih.gov/ij/) by measuring the mean alveolar fluorescence intensity (MFI) from randomly selected ROIs. The MFI was determined from 4-5 ROIs of each patient sample by setting a same intensity threshold for CC and FC-treated hPCLS.

Quantification of epithelial cell types and states, including KRT17+/KRT5-basaloid cells was achieved by using the “labeled-based object classification by surface components” feature of IMARIS 9.6.0 software (Bitplane). Single cell segmentations and object classification was done with the ImageJ-LABKIT extension^93^, which performs a machine-learning based classification built of the distinctive shape of a cell type of interest. Multiple classes were added and defined in order to label and determine the number of cells under a particular combination of fluorescent signals. For the final analysis, the entire area of the hPCLS punches in the FFPE sections was analyzed for KRT17, KRT5 and SFTPC positive cells, as well as DAPI for total cell counts per punched area, which was used for normalization. In sum 130.636 single cells from eight different patients (n=8) were analyzed.

Quantification of epithelial cell types and states in time-resolved perturbations (FC) after 12h, 24h, and 48h was accomplished by analyzing FFPE sections from punched hPCLS, which were immunostained for KRT17, KRT5 and SFTPC along with DAPI for cell nuclei. Cells were quantified and subsequently classified (as depicted in Fig.3g) according to their morphology and protein expression phenotypes, that is, cuboidal SFTPC+/KRT5-/KRT17-; cuboidal SFTPC+/KRT5-/KRT17+; elongating SFTPC+/KRT5-/KRT17+; elongated SFTPC-/KRT5-/KRT17+ at various time-points (12h, 24h, 48h). In sum 120 ROIs were assessed. Each ROI measured 500 µm x 500 µm. In total 1,136 single cells from four different patients (n=4) were analyzed.

### MicroCT based staging of human IPF patient tissue

The lung processing protocol is described extensively elsewhere^35, 94^. Lungs were air-inflated upon total lung capacity after explantation and immediately frozen in the fumes of liquid nitrogen. After conventional scanning using computed tomography, the lungs were processed into cores (2cm height, 1.5 cm diameter). These cores were scanned using microCT (Skyscan, Belgium) while frozen. Afterwards, disease extent was calculated using the microCT-derived parameter Surface Density (SfD), which was shown to negatively correlate with Ashcroft score^25^. Using SfD cut-offs, the samples were classified in mild (i.e. IPF1), moderate (i.e. IPF2) and severe (i.e. IPF3) fibrosis. In case microCT data was not available, samples were classified according to simplified Ashcroft scoring (one scoring for the entire core) in mild (Ashcroft 1-4), moderate (Ashcroft 5-6) and severe (Ashcroft 7-8).

### Immunofluorescence staining, microscopy and quantification of human IPF patient and donor tissues

Stainings of human IPF patient and control tissues were performed following the same protocol as used for the FFPE sections of hPCLS (see above). In brief, lung tissues derived from staged IPF and control tissues were fixed in 10% neutral-buffered formalin (Sigma) overnight at 4°C, followed by dehydration, paraffin infiltration and sectioning into 4 micron thick FFPE sections. After drying, FFPE sections were deparaffinized, rehydrated, subjected to antigen retrieval with 10 mM citric acid buffer and blocked with 10% normal donkey serum for 1 hour at RT. The same primary antibodies were used as for the *ex vivo* PCLS FFPE slices with optimized concentrations for FFPE sections from patient tissues (CLDN5 1:100; PLVAP 1:300; KRT5 1:1000, KRT17 1:200; SFTPC 1:50,PDPN 1:200, R&D, AF3670). After incubation at 4°C overnight, FFPE sections were incubated with the appropriate secondary antibodies (donkey anti-sheep Alexa Fluor 568, A20042, donkey anti-mouse Alexa Fluor 488, A21202, 1:250; donkey anti-rabbit Alexa Fluor 647, A31573, 1:250, goat anti-chicken Alexa Fluor 568, A11041, 1:250; all from Invitrogen) for 2h at RT followed by nuclear counterstain with DAPI (10 min, RT). FInally, unspecific staining due to tissue autofluorescence was blocked using the Vectastain TrueVIEW autofluorescence blocking kit (Vector Laboratories). Whole slide imaging was performed using the Axioscan 7 slidescanner (Zeiss).

Percentages of epithelial cell types and states were quantified in four randomly selected ROIs from each patient analogously to the *ex vivo* hPCLS data using IMARIS 9.6.0 software (see above).

PLVAP signals were quantified in ImageJ and multiple regions of interest (ROIs, n=4-5) from control tissues and microCT staged IPF tissues were randomly selected to measure the total positive areas of all EC markers. The area fraction of PLVAP+ vessels was then determined by dividing the positive area to the total area of each ROIs.

Quantification of the appearance and abundance of PLVAP+ ectopic vessels in alveolar septa (AS), was performed in ImageJ (https://imagej.nih.gov/ij/). Briefly, we randomly selected around 120 alveolar septa (60 of each condition) from 5 control and 4 IPF stage-1 tissues, and the alveolar septal area and length were determined based on PDPN expression. The main steps included (1) thresholding of the PDPN positive area, (2) conversion into a binary image, (3) filling of holes, and (4) determining the size of total alveolar septal area after producing a solid AS mask. Additionally, the alveolar length was measured by drawing a central line for each AS so that the mean AS thickness could be computed by dividing the total AS area to the AS length. Finally, The PLVAP+ area was calculated by setting intensity thresholds for all AS samples, and normalized to the individual AS length.

To investigate the relationship of AS thickness and abundance of PLVAP+ vessels, Pearson correlation analysis was performed between the relative PLVAP+ vessel area/AS length and the AS thickness.

### Iterative indirect immunofluorescence imaging

For *in situ* validation of basaloid to myofibroblast crosstalk, we performed iterative indirect immunofluorescence imaging (4i)^64^ on control and microCT staged IPF tissues staining for epithelial (KRT17, KRT5, KRT8, SFTPC, PDPN) and mesenchymal (TNC, aSMA, DESMIN) markers.

As for the conventional IF stainings, for 4i, FFPE sections were first deparaffinized, dehydrated and subjected to heat-mediated antigen retrieval with 1X R-universal antigen retrieval buffer (Aptum). Next, nonspecific antibody binding was blocked for 1 hour at RT with 1% BSA supplemented with 150 mM maleimide. Afterwards, primary antibodies were added and incubated overnight at 4°C. After washing, secondary antibodies were added and incubated for 2h at RT followed by DAPI staining and autofluorescence blocking. To avoid photocrosslinking during microscopy, sections were immersed in imaging buffer containing 700 mM N-Acetyl-cysteine and 20% 1 M HEPES (pH=7.4) and subjected to whole slide imaging using the Axioscan 7 slidescanner (Zeiss).

After elution of antibodies using a buffer (pH=2.5) containing 0.5M L-Glycine, 3M Urea, 3M guanidinium chloride, and 70mM TCEP-HCl, antibody staining (without deparaffinization, dehydration and antigen retrieval) and imaging as described above were repeated until the desired plexity was achieved (5 repeats).

The following primary (1) and secondary antibodies (2) were used: (1) PDPN 1:100, R&D, AF3670; KRT5 1:800, biolegend, 905901; KRT17 1:200, Sigma, HPA000453; SFTPC 1:150, Sigma, HPA010928; KRT8 1:200, University of Iowa Hybridoma Bank; DESMIN 1:100, Santa Cruz, sc-23879, aSMA, 1:1500, Sigma, A5228; TNC, 1:100, abcam, ab108930; (2) donkey anti-rabbit Alexa Fluor 488, A21206; donkey anti-sheep Alexa Fluor 568, A20042; donkey anti-mouse Alexa Fluor 647, A31571; donkey anti-mouse Alexa Fluor 568, A10037; donkey anti-goat Alexa Fluor 488, A11055; donkey anti-rabbit Alexa Fluor 568, A10042; donkey anti-rat Alexa Fluor 647, A48272; donkey anti-rabbit Alexa Fluor 647, A31573; donkey anti-sheep Alexa Fluor 488, A11015; (all from Invitrogen at 1:250); donkey anti-chicken AF488, Jackson, 703-545-155, 1:250.

For *ex vivo* validation in FFPE sections of hPCLS treated with CC and FC cocktails for 48 hours or 6 days, a subset of selected epithelial (KRT17 1:400, SFTPC 1:50, PDPN 1:100) and mesenchymal (aSMA 1:1500, TNC 1:100, DESMIN 1:100) was stained and imaged using the same protocol.

Alignment of microscopy images of different cycles was performed by overlaying selected regions of interest and correcting the slight shifts in X and Y position that occurred between the acquisition cycles using the individual DAPI signals as reference in ZEN Blue software (Zeiss).

## List of Supplementary Materials

**Supplementary table S1:** Baseline characteristics of included human IPF and control samples from Leuven

**Supplementary table S2**: Marker genes *ex vivo* (meta cell type level)

**Supplementary table S3**: Marker genes *ex vivo* (cell type level)

**Supplementary table S4**: Marker genes *in vivo* (meta cell type level)

**Supplementary table S5**: Marker genes *in vivo* (cell type level)

**Supplementary table S6:** Gene expression changes *in vivo* and *ex vivo* (in PF, after FC, FC+CMP4, and FC+Nintedanib treatment) for each meta cell type

**Supplementary table S7**: Gene expression changes of conserved cell-circuit cell states (basaloid, ectopic ECs, myofibroblasts, pericytes, SPP1+ Macrophages) after CMP4 and Nintedanib treatment

## Data availability

Raw 10X Genomics data generated for this study will be made available with limited access via the Genome-Phenome Archive (EGA). Processed count tables are available on Zenodo (10.5281/zenodo.7537493). The original, publicly available single-cell RNA-seq datasets used to generate the integrated cell atlas of Pulmonary Fibrosis can be accessed under the following GEO accession numbers: GSE136831 (New Haven cohort), GSE122960 (Chicago cohort), GSE135893 (Nashville cohort), and under https://github.com/theislab/2020_Mayr (Munich cohort). The original, publicly available bulk RNA-seq data from microCT based staged IPF tissues re-analysed in this study can be accessed under the following GEO accession number: GSE124685.

## Code availability

Code and Jupyter notebooks that were used to analyze the data alongside the .yml files of the respective conda environments used are deposited at GitHub: https://github.com/niklaslang/PCLS_perturbations

## Supplementary Figures

**Supplementary Figure S1.**
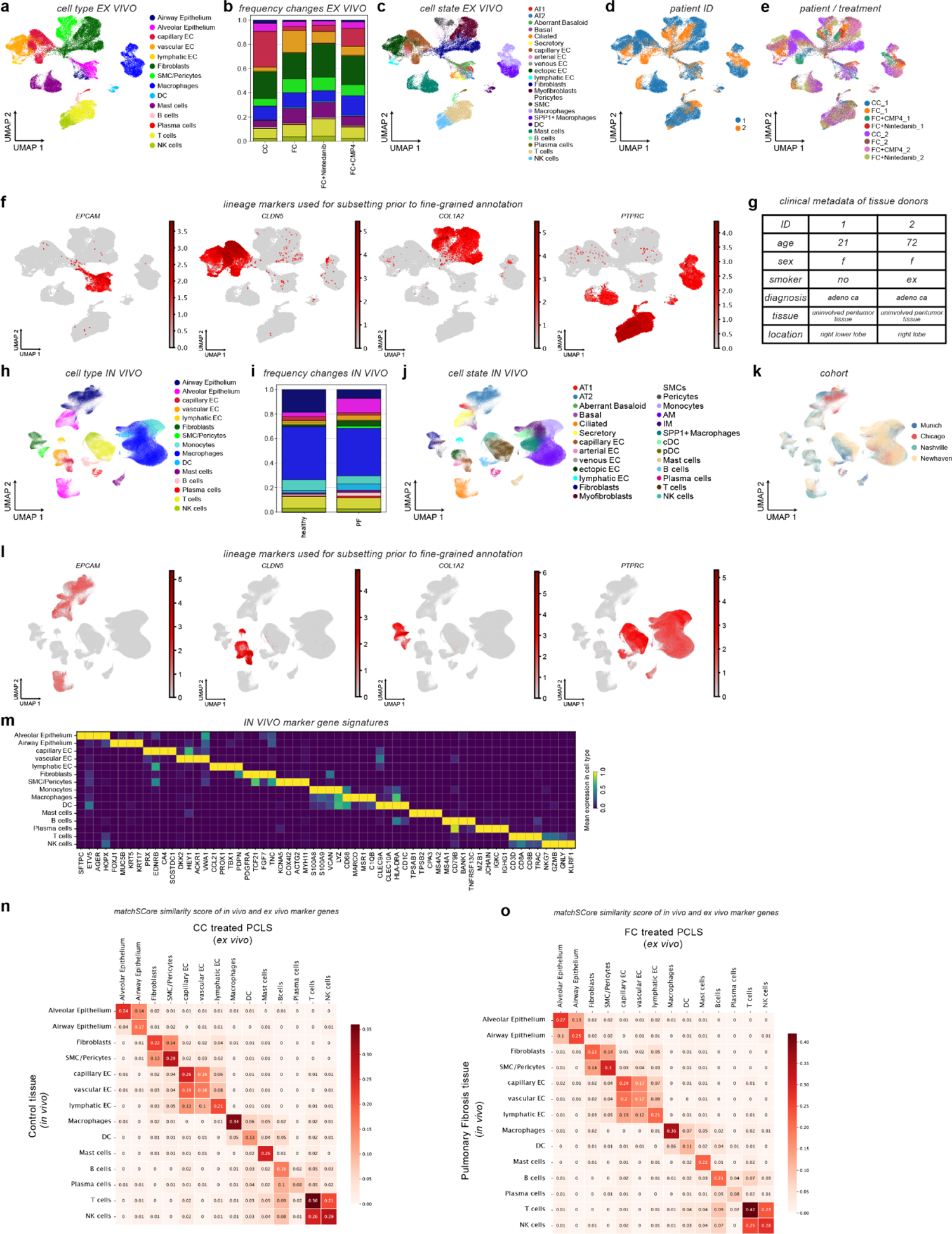
*Ex vivo* hPCLS recapitulates early-stage events in pulmonary fibrosis (corresponding to Figure 1). (a) UMAP embedding of 63,581 single cells from hPCLS color coded by meta cell type identity. (b) Stacked bar plots show relative cell type frequency changes with regards to treatment. (c-e) UMAP embedding of hPCLS data color coded by cell state (c), tissue donor (d) and tissue donor/treatment (e), respectively. (f) UMAP feature plots show relative expression of lineage markers EPCAM+ (epithelial cells), CLDN5*+* (endothelial cells), COL1A2*+* (stromal cells), and PTPRC*+* (CD45, immune cells) cells in the *ex vivo* hPCLS data. (g) Clinical metadata of hPCLS tissue donors. (h) UMAP embedding of 481,788 single cells from the integrated multi-cohort pulmonary fibrosis cell atlas, color coded by meta cell type identity. (i) Stacked bar plot of relative cell type frequency changes. (j,k) UMAP embedding of the integrated pulmonary fibrosis cell atlas color coded by cell state (j) and cohort (k), respectively. (l) UMAP feature plots UMAP feature plots showing relative expression of lineage markers EPCAM+ (epithelial cells), CLDN5+ (endothelial cells), COL1A2+ (stromal cells), and PTPRC+ (CD45, immune cells) in the *in vivo* integrated pulmonary fibrosis cell atlas (m) *In vivo* marker genes signatures of meta-cell type identities. (n,o) matchSCore comparison of *ex vivo* against *in vivo* marker gene signatures from Control Cocktail (CC) treated hPCLS (d6) and control lungs (n) as well as Fibrotic Cocktail (FC) treated hPCLS (d6) and pulmonary fibrosis lungs (o).

**Supplementary Figure S2.**
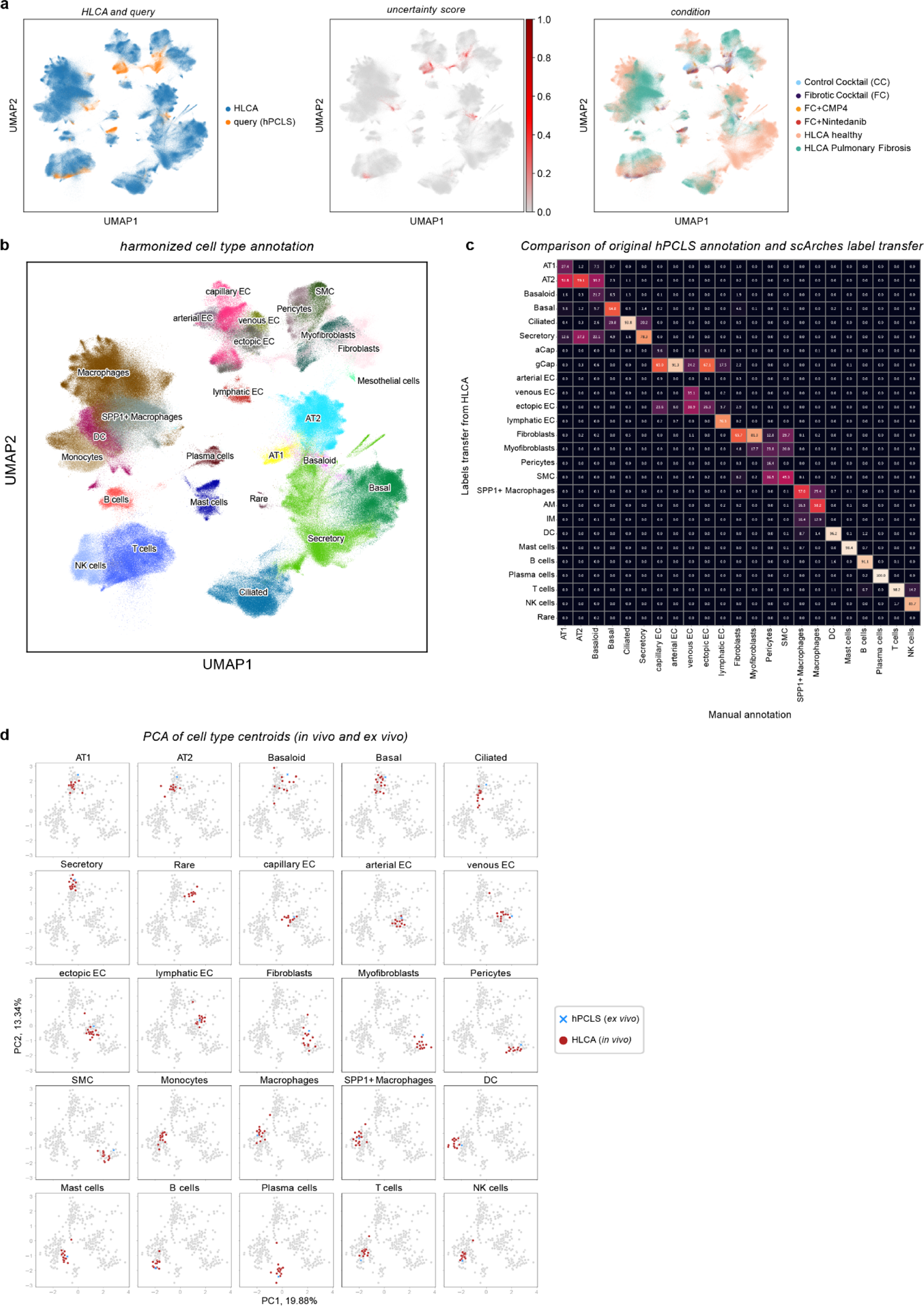
Query-to-reference mapping of hPCLS data to the HLCA (corresponding to Figure 1). (**a**) UMAP embeddings of 922,332 single cells from the PF-extended HLCA (*in vivo* reference) and 63,581 single cells from hPCLS (*ex vivo* query) after scArches mapping color coded by reference/query (left), label transfer uncertainty score (center), and condition (right). (**b**) UMAP embedding of 922,332 single cells from the HLCA (*in vivo* reference) and 63,581 single cells from hPCLS (*ex vivo* query) after scArches mapping color coded by the final, harmonized cell type annotation labels. (**c**) Comparison of original hPCLS annotation and scArches label transfer. Heatmap visualizes percentage overlap between original hPCLS annotation (columns) and scArches label transfer (rows). (**d**) PCA plot of per sample cell type centroids in the latent embedding space demonstrating transcriptional proximity of cell identities from hPCLS (*ex vivo* query) to the PF-extended HLCA (*in vivo* reference; healthy and PF).

**Supplementary Figure S3.**
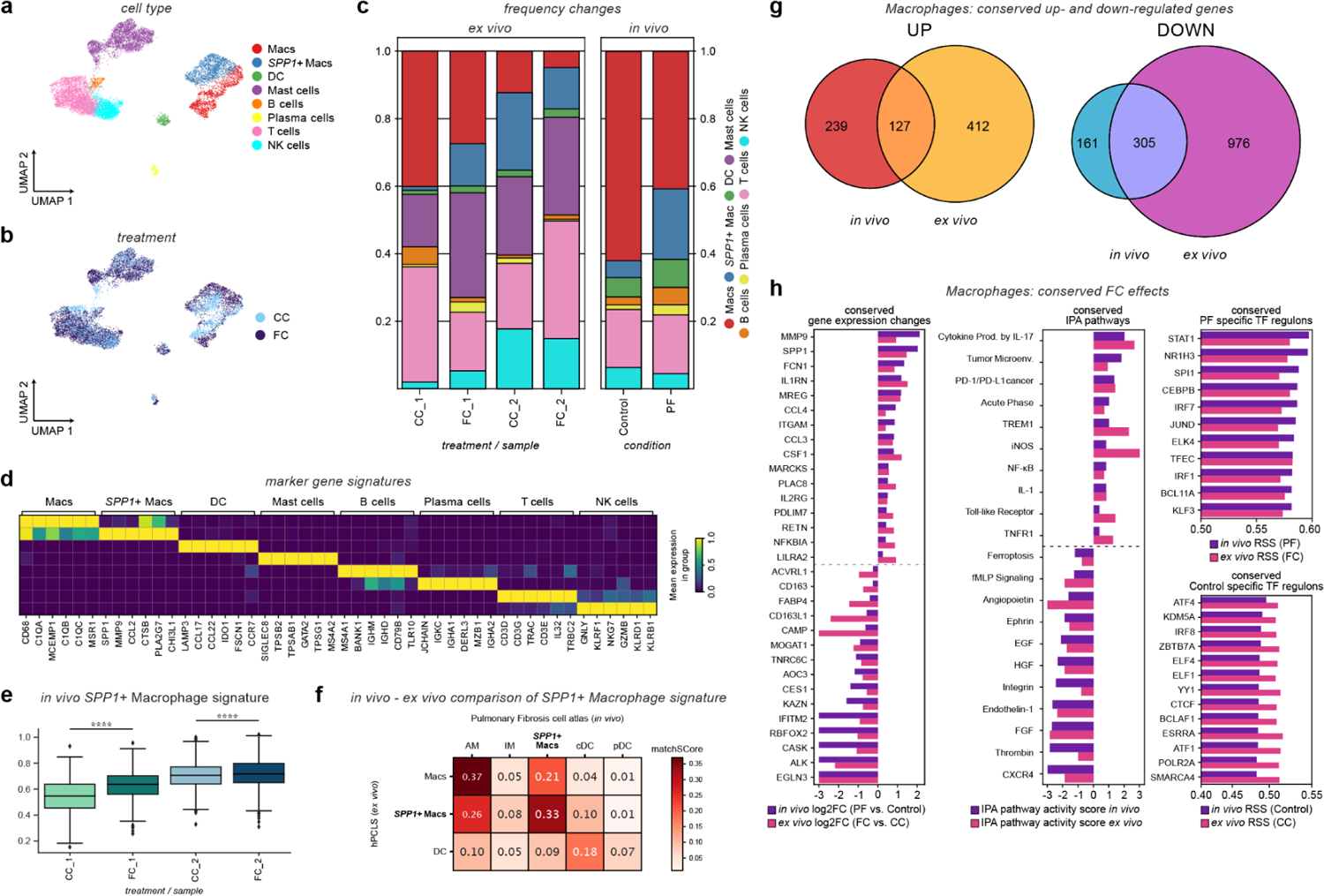
Ex vivo SPP1+ macrophage state with similarities to PF associated macrophages. **(a,b)** UMAP embedding of 13,239 single PTPRC+ immune cells from FC and CC treated hPCLS, color coded by cell type identity **(a)** and treatment **(b)**. **(c)** Cell type proportion analysis of immune cells in FC treated hPCLS (*ex vivo*, left) and Pulmonary Fibrosis (PF) (*in vivo*, right): Stacked bar plot shows the relative frequency of each cell type with regards to treatment and tissue donor. **(d)** Ex vivo marker gene signatures of immune cells. The heatmap shows the average scaled expression in each cell type. **(e)** Boxplot of *in vivo* SPP1+ macrophage gene signature (derived from the *in vivo* reference Pulmonary Fibrosis cell atlas) score after treatment with FC with regards to treatment and tissue donor. Mann-Whitney-Wilcoxon test, ****p ≤ 0.0001. **(f)** matchSCore comparison of *ex vivo* against *in vivo* marker genes of all mononuclear phagocytic cells, highlighting highest transcriptional similarity of *ex vivo* SPP1+ Macrophages with *in vivo* SPP1+ Macrophages followed by *in vivo* Alveolar Macrophages. **(g)** Quantitative side-by-side comparison of genes differentially expressed in Macrophages *in vivo* and *ex vivo*. The venn diagram illustrates the intersection of genes that are uniformly upregulated in Macrophages both *in vivo* and *ex vivo*. **(h)** Qualitative analysis of conserved responses to FC on the gene, pathway and Transcription Factor (TF) regulon level in Macrophages. The bar plots illustrate the uniform behavior of conserved genes (log2FC), PF- and health-specific TF regulons (Regulon Specificity Score - RSS), and IPA pathways (IPA pathway activity score) in Macrophages. Abbreviations: Macs: Macrophages, AM: Alveolar Macrophages, IM: Interstitial Macrophages, cDC: conventional Dendritic cells, pDC: plasmacytoid Dendritic cells.

**Supplementary Figure S4.**
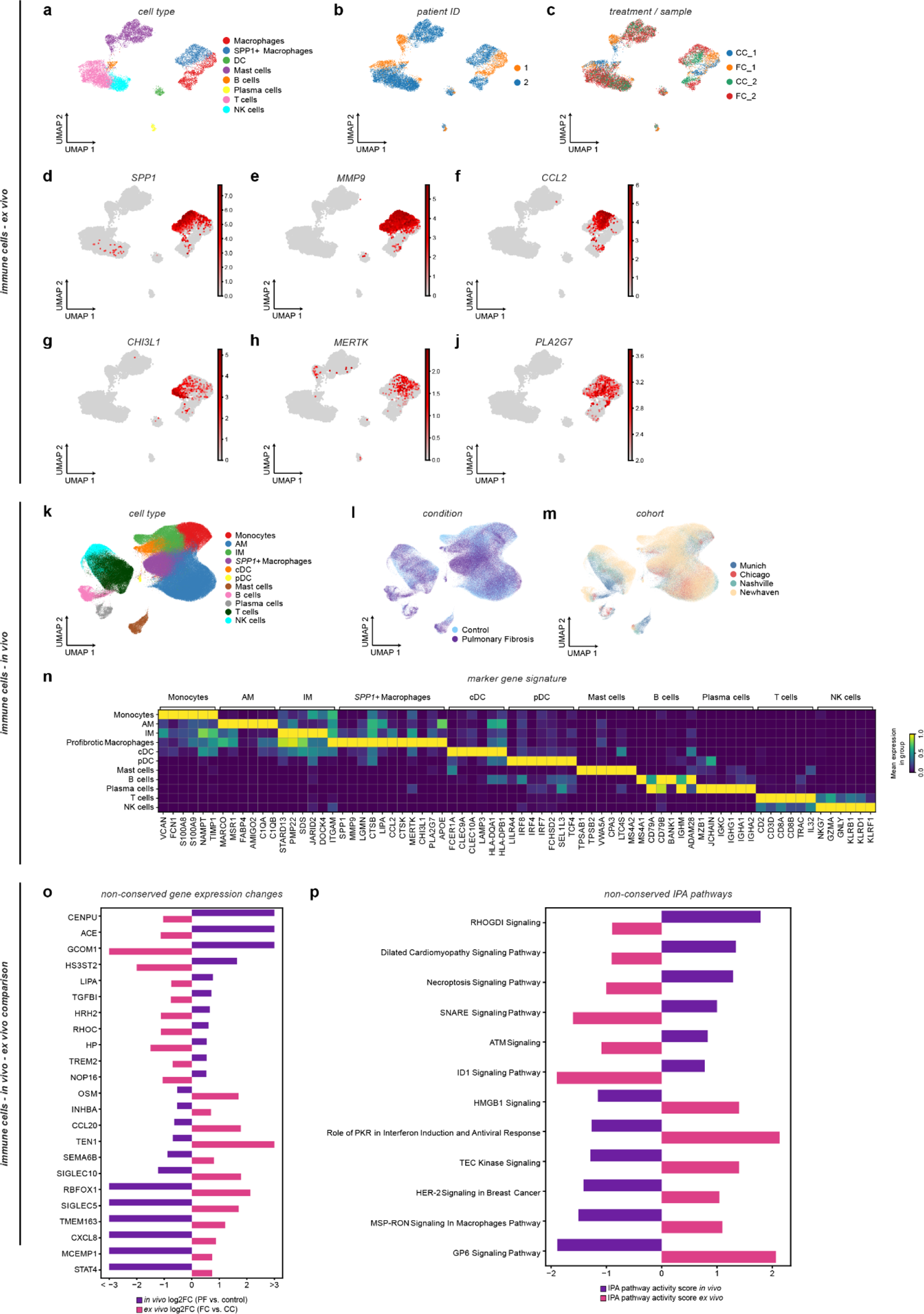
Myeloid phenotypes in FC treated hPCLS. (corresponding to Supp. Fig. S3). **(a-c)** UMAP embedding of 13,239 single PTPRC+ immune cells from FC and CC treated hPCLS, color coded by cell type **(a)**, tissue donor **(b)**, and treatment / sample **(c)**. **(d-j)** UMAP feature plots illustrating relative expression of SPP1+ macrophages marker gene signature. **(k-m)** UMAP embeddings of 328,410 single PTPRC+ immune cells from the *in vivo* pulmonary fibrosis cell atlas, color coded by cell type **(k)**, disease status **(l)**, and cohort **(m)**. **(n)** *In vivo* marker gene signatures. The heatmap shows the average scaled expression of markers in each immune cell type. **(o,p)** Qualitative analysis of non-conserved responses to FC on the gene **(o)** and pathway **(p)** level in macrophages. The bar plots illustrate the diverging behavior of non-conserved genes (log2FC) **(o)**, and IPA pathways (IPA pathway activity score) **(p)** in macrophages.

**Supplementary Figure S5.**
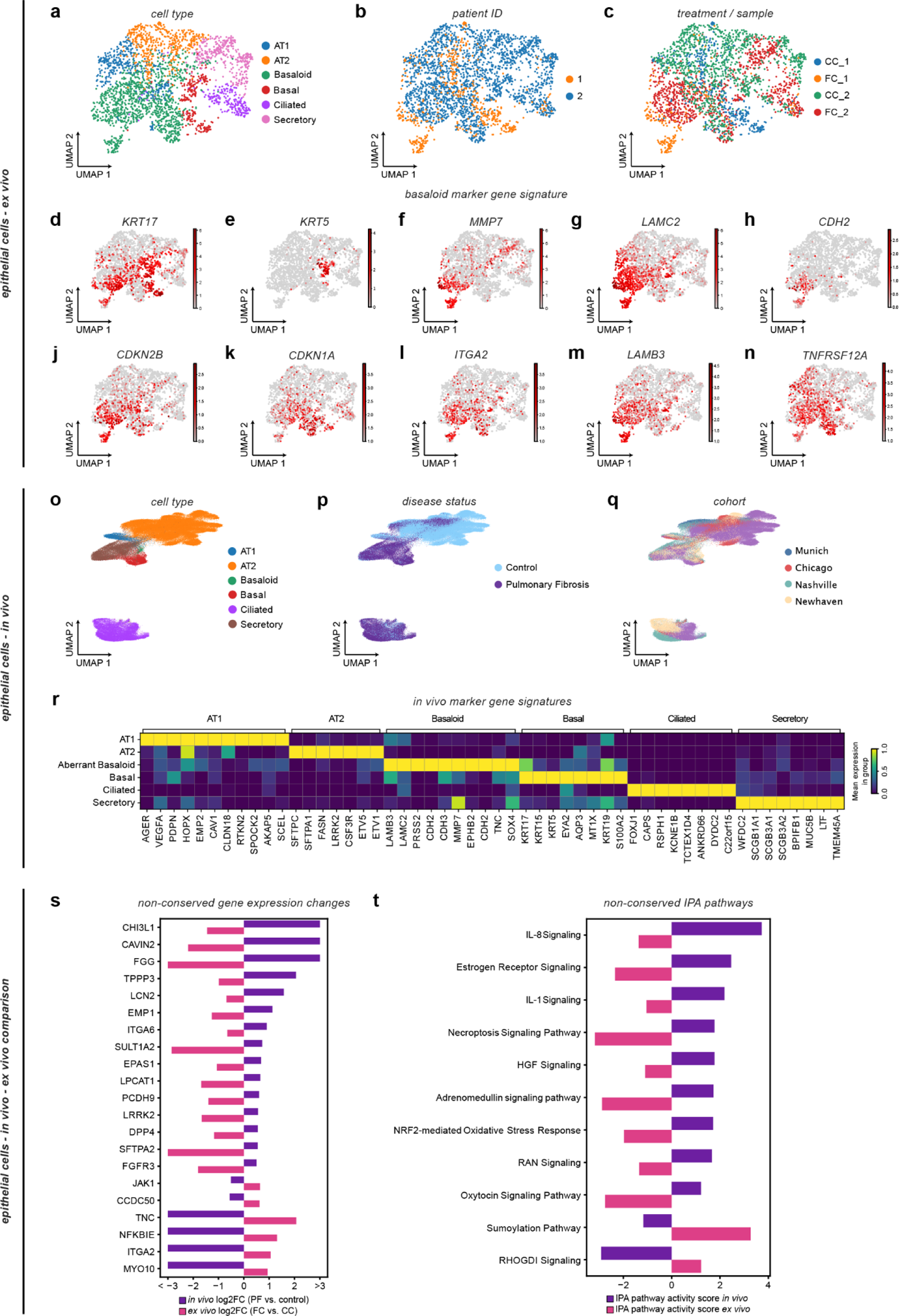
Induction of KRT17+/KRT5-basaloid cells in *ex vivo* hPCLS (corresponding to Figure 2). **(a-c)** UMAP embedding of 2,741 single EPCAM+ epithelial cells from FC and CC treated hPCLS, color coded by cell type **(a)**, tissue donor **(b)**, and treatment / sample **(c)**. **(d-n)** UMAP feature plots illustrating relative expression of KRT17+/KRT5-basaloid marker gene signature. **(o)** DC embeddings of trajectory inference analysis on AT2 cells, basaloid cells, and basal cells color coded (from left to right) cell type overlaid by RNA velocity, diffusion pseudotime, condition, relative expression of AT2 marker SFTPC, relative expression of shared basal and basaloid cell marker KRT17 and relative expression of basaloid marker MMP7. **(p-r)** UMAP embeddings of 181,453 single EPCAM+ epithelial cells from the *in vivo* pulmonary fibrosis cell atlas, color coded by cell type **(p)**, disease status **(q)**, and cohort **(r)**. **(q)** *In vivo* marker gene signatures. The heatmap shows the scaled average expression of markers in each epithelial cell type. **(t,u)** Qualitative analysis of non-conserved responses to FC on the gene **(t)** and pathway **(u)** level in alveolar epithelial cells. The bar plots illustrate the diverging behavior of non-conserved genes (log2FC) **(t)**, and IPA pathways (IPA pathway activity score) **(u)** in alveolar epithelial cells.

**Supplementary Figure S6.**
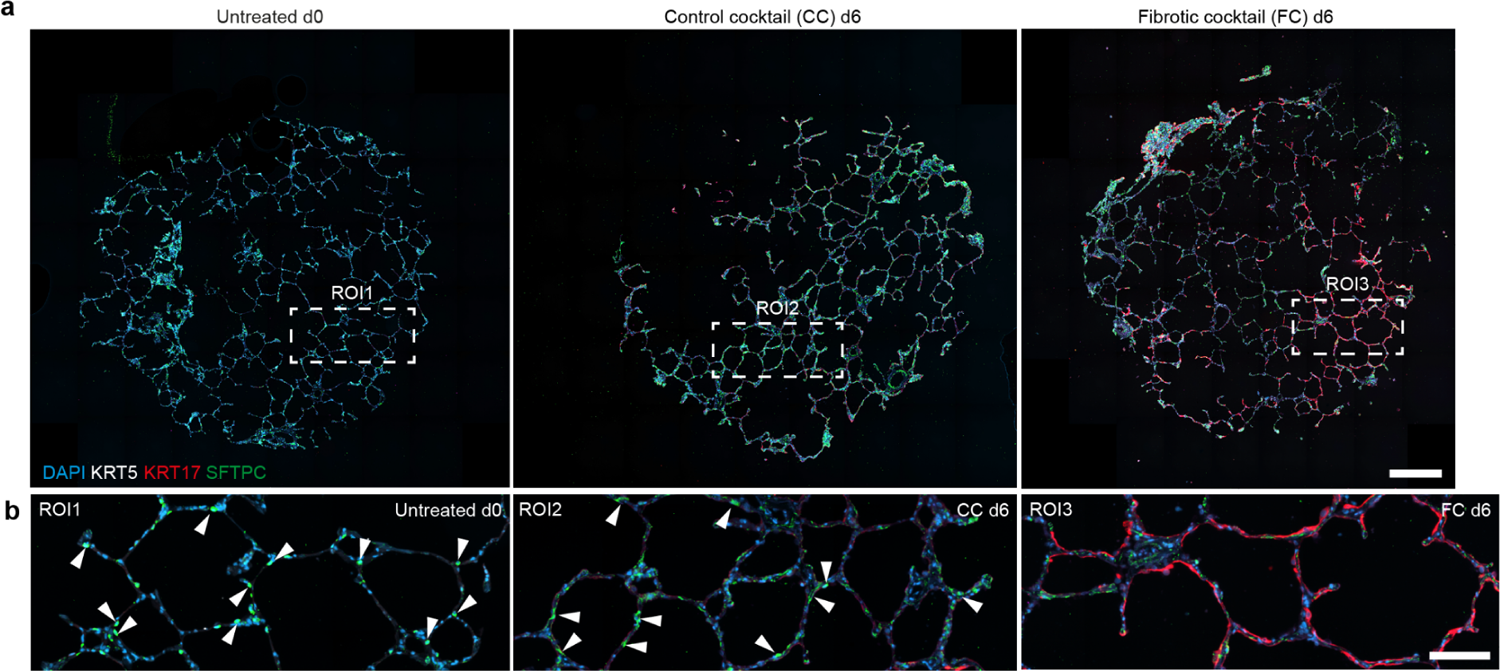
FC treatment induces the appearance of KRT17+ cells (corresponding to Figure 3). **(a)** IF of FFPE-punches from hPCLS shown at day 0, and day 6 following CC and FC treatment. Scale bar = 500 µm. **(b)** Arrowheads indicate cuboidal shaped SFTPC+ AT2 cells (in green, ROI1). SFTPC+ cells (arrowheads) are still present after day 6 of CC treatment (in green, ROI2). On day 6 after FC treatment, SFTPC+ cells mostly disappeared from areas of high KRT17+ expression (in red, ROI3). Cell nuclei were stained with DAPI (blue). Scale bar = 50 µm.

**Supplementary Figure S7.**
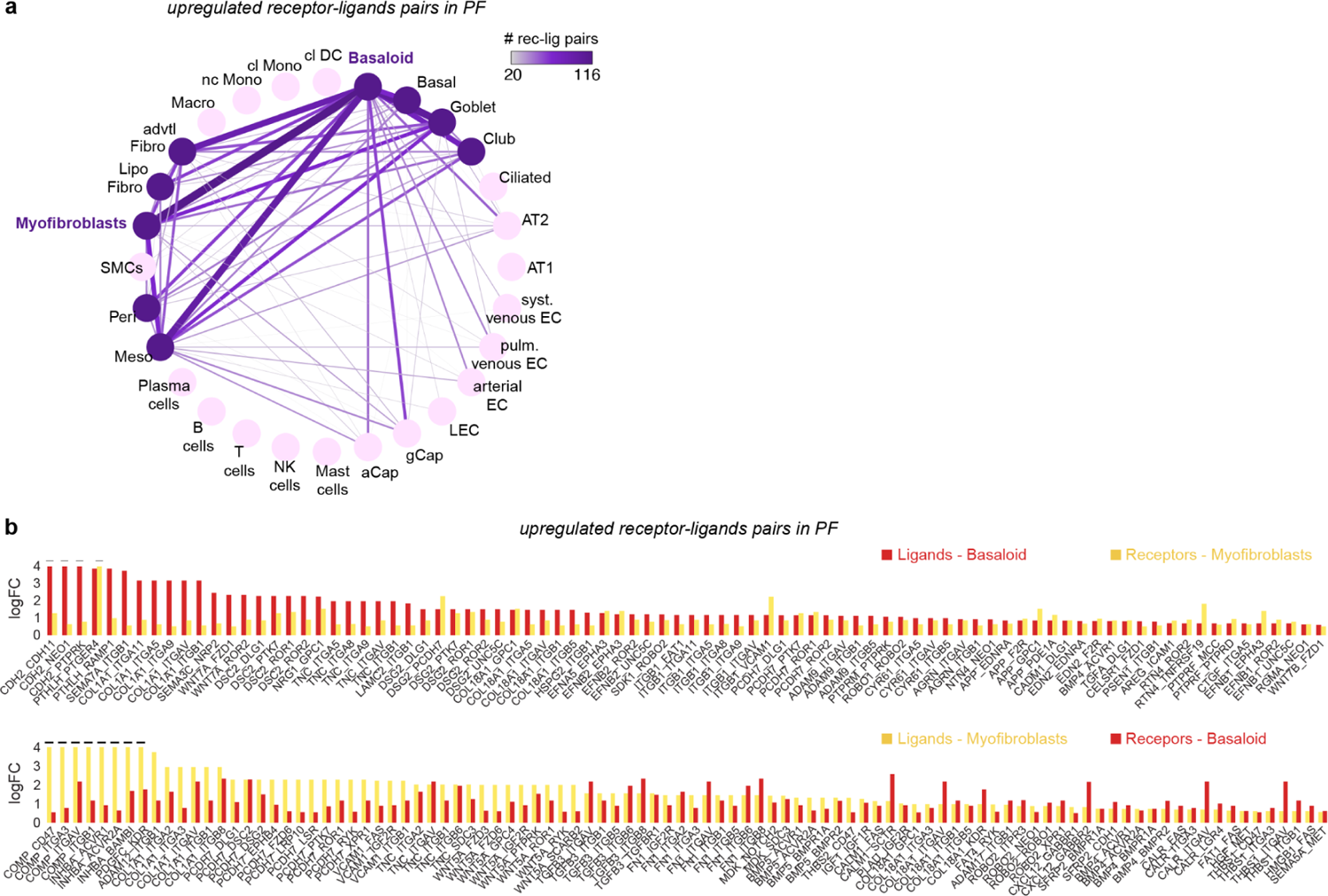
Cell-cell communication between basaloid cells and myofibroblasts. **(a)** Number upregulated of receptor-ligand pairs in the integrated multi-cohort PF cell atlas. **(b)** Log2 foldchanges of between basaloid cells and myofibroblasts.

**Supplementary Figure S8.**
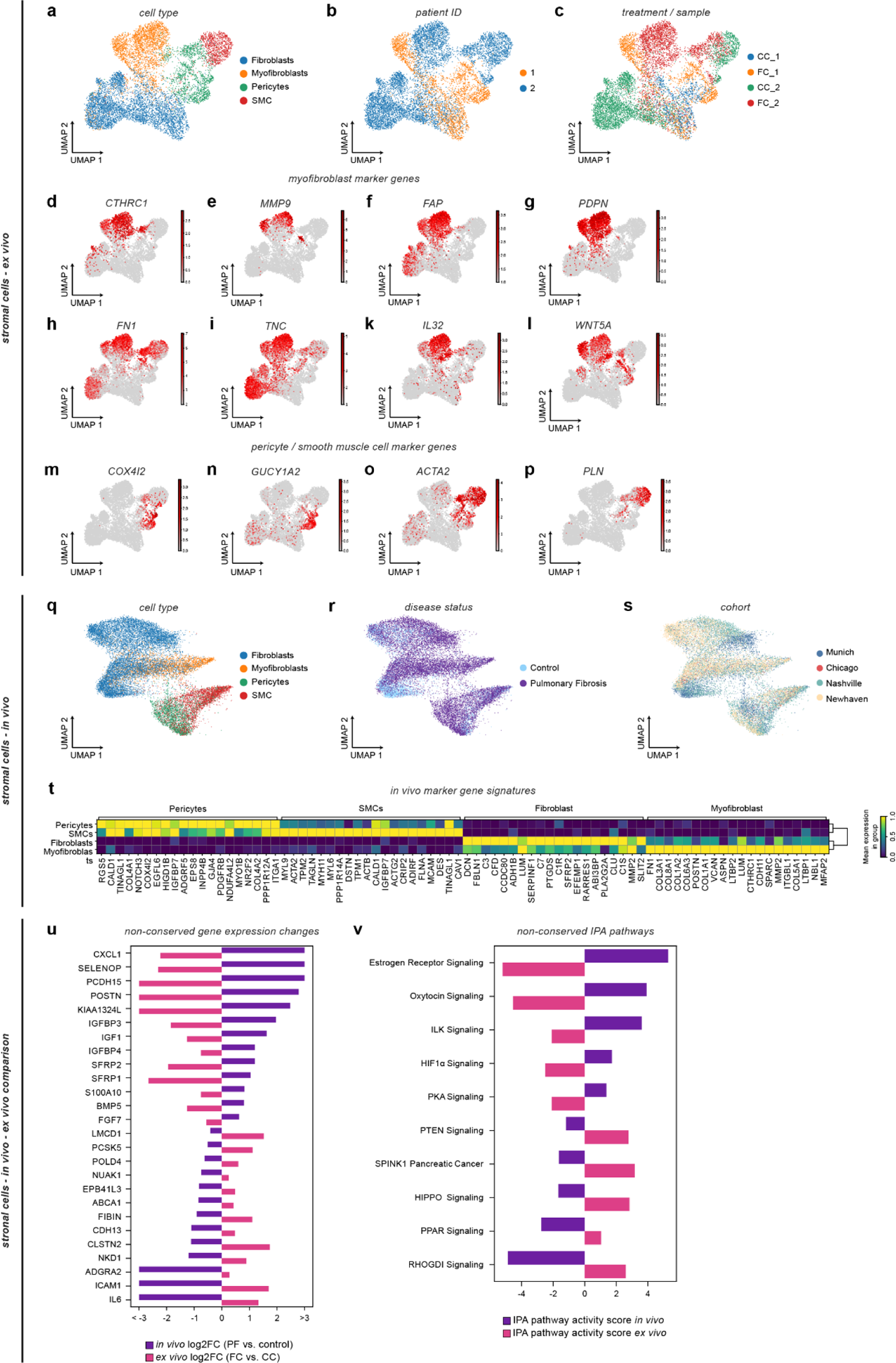
Induction of CTHRC1+ myofibroblasts in *ex vivo* hPCLS (corresponding to Figure 4). **(a-c)** UMAP embedding of 9,721 single stromal cells from FC and CC treated hPCLS, color coded by cell type identity **(a)**, tissue donor **(b)**, and treatment/sample **(c)**. **(d-l)** UMAP feature plots illustrating relative expression of CTHRC1+ myofibroblast marker gene signature. **(m-p)** UMAP feature plots illustrating relative expression of pericytes and smooth muscle cells marker genes. **(q-s)** UMAP embeddings of 19,335 single COL1A2+ stromal cells from the *in vivo* pulmonary fibrosis cell atlas, color coded by cell type **(q)**, disease status **(r)**, and cohort **(s)**. **(t)** *In vivo* marker gene signatures. The heatmap shows the average scaled expression of markers in each stromal cell type. **(u,v)** Qualitative analysis of non-conserved responses to FC on the gene **(u)** and pathway **(v)** level in (myo)fibroblasts. The bar plots illustrate the diverging behavior of non-conserved genes (log2FC) **(u)**, and IPA pathways (IPA pathway activity score) **(v)** in (myo)fibroblasts.

**Supplementary Figure S9.**
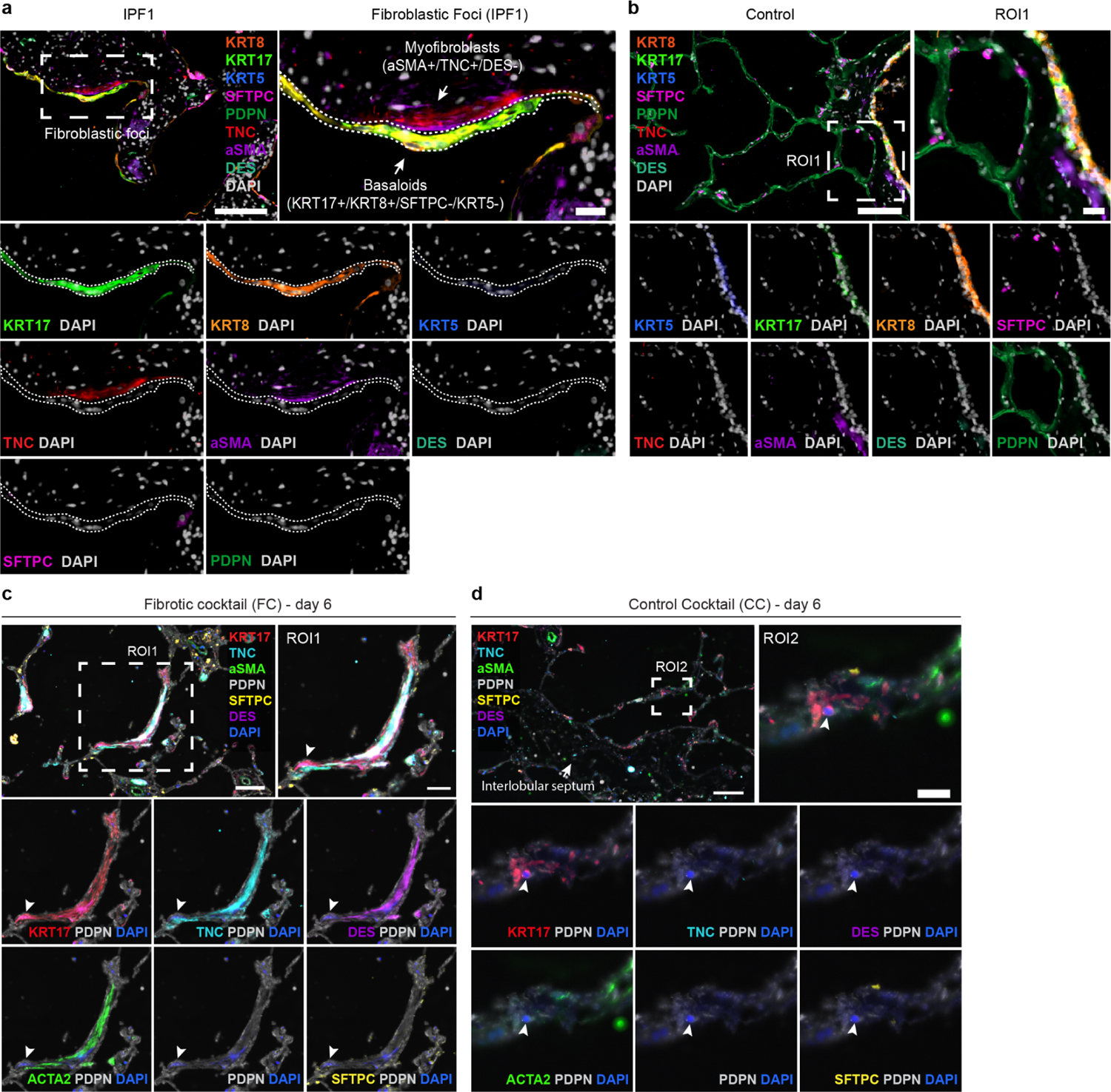
Myofibroblast-Basaloid cell interactions are absent in control lungs yet define fibroblast foci in early-stage IPF and are recapitulated in the hPCLS model upon fibrotic cocktail treatment (corresponding to Figure 4). **(a)** Additional representative indirect iterative IF image (4i) of eight different epithelial (KRT17, KRT5, KRT8, SFTPC, PDPN) and stromal (TNC, aSMA, DES) cell markers demonstrating the interaction of elongated basaloid (SFTPC-/KRT17+/KRT8+/KRT5-) cells (white dashed contour) and myofibroblasts (TNC+/aSMA+/DES-) in fibroblastic foci of early-stage IPF (IPF1). Scale bars = 100 µm (overview image) and 20 µm (enlarged view). **(b)** Basaloid cells and myofibroblasts are absent in control lungs. **(c)** The interaction of SFTPC-/KRT17+ cells (white arrowheads) and TNC+/aSMA-/DES-(myo-)fibroblasts is also recapitulated *ex vivo* in FC-treated hPCLS at d6. In addition, an interaction of KRT17+/SFTPC-cells with TNC+/aSMA+/DES+ cells in the thickened alveolar septum has been observed. Scale bars = 100 µm (overview image) and 50 µm (enlarged view). **(d)** No interaction of KRT17+ cells and (myo-)fibroblasts are found in CC-treated hPCLS (d6). Scale bars = 100 µm (overview image) and 20 µm (enlarged view).

**Supplementary Figure S10:**
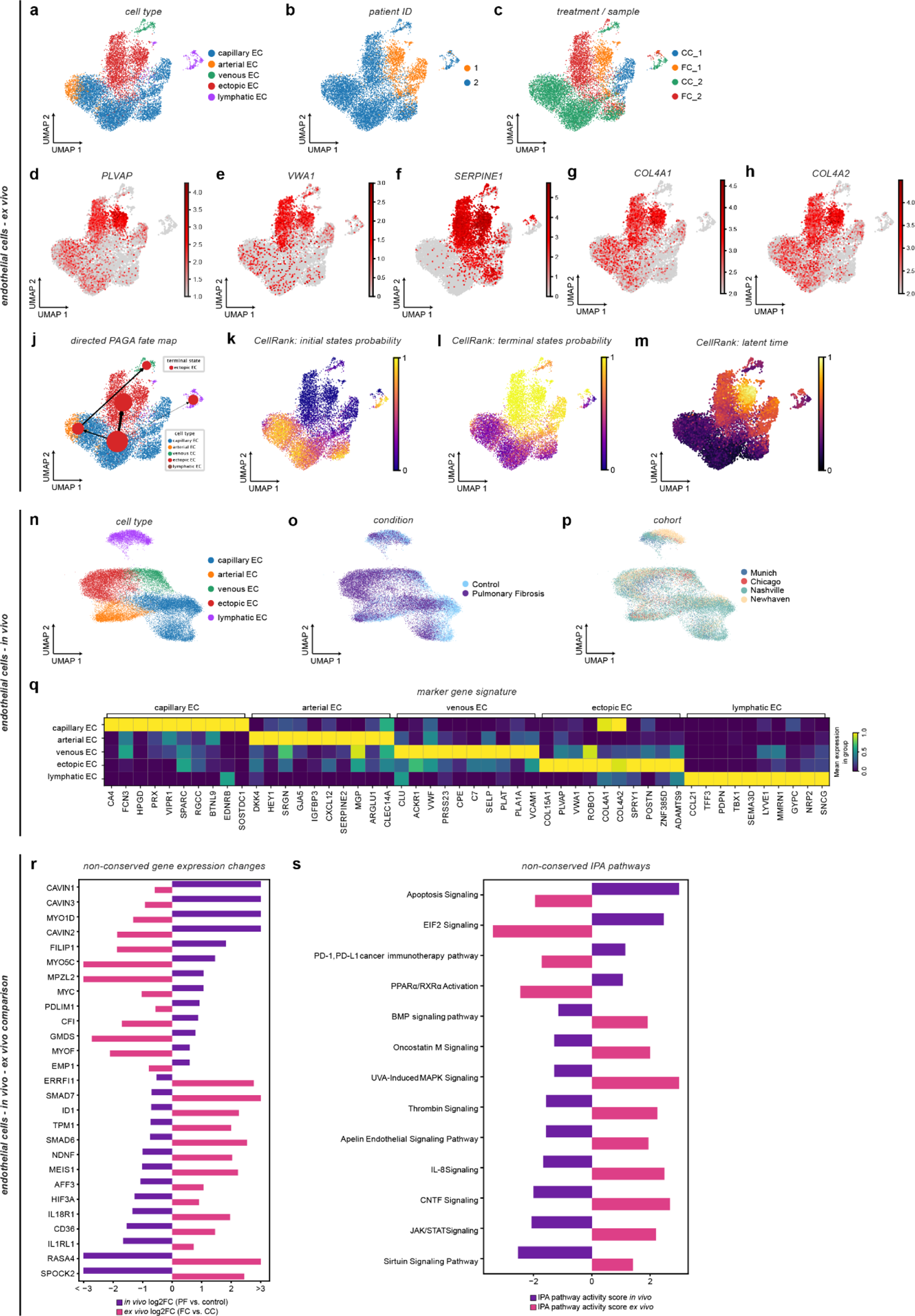
Induction of VWA1+/PLVAP+ ectopic ECs in *ex vivo* hPCLS (corresponding to Figure 5). **(a-c)** UMAP embedding of 10,418 single CLDN5+ epithelial cells from FC and CC treated hPCLS, color coded by cell type **(a)**, tissue donor **(b)**, and treatment / sample **(c)**. **(d-h)** UMAP feature plots illustrating relative expression of VWA1+/PLVAP+ ectopic EC marker gene signature. **(j-m)** UMAP embeddings of trajectory inference analysis on endothelial cells, color coded by cell type and overlaid by estimated cell type connectivity as measure by partition-based graph abstraction (PAGA) **(j)**, CellRank initial states probability **(k)**, CellRank terminal states probability **(l)**, and CellRank latent time **(m)**. **(n-p)** UMAP embeddings of 29,534 single CLDN5+ immune cells from the *in vivo* pulmonary fibrosis cell atlas, color coded by cell type **(n)**, disease status **(o)**, and cohort **(p)**. **(q)** *In vivo* marker gene signatures. The heatmap shows the average scaled expression of markers in each endothelial cell type. **(r,s)** Qualitative analysis of non-conserved responses to FC on the gene **(r)** and pathway **(s)** level in vascular endothelial cells. The bar plots illustrate the diverging behavior of non-conserved genes (log2FC) **(r)**, and IPA pathways (IPA pathway activity score) **(s)** in vascular endothelial cells.

**Supplementary Figure S11.**
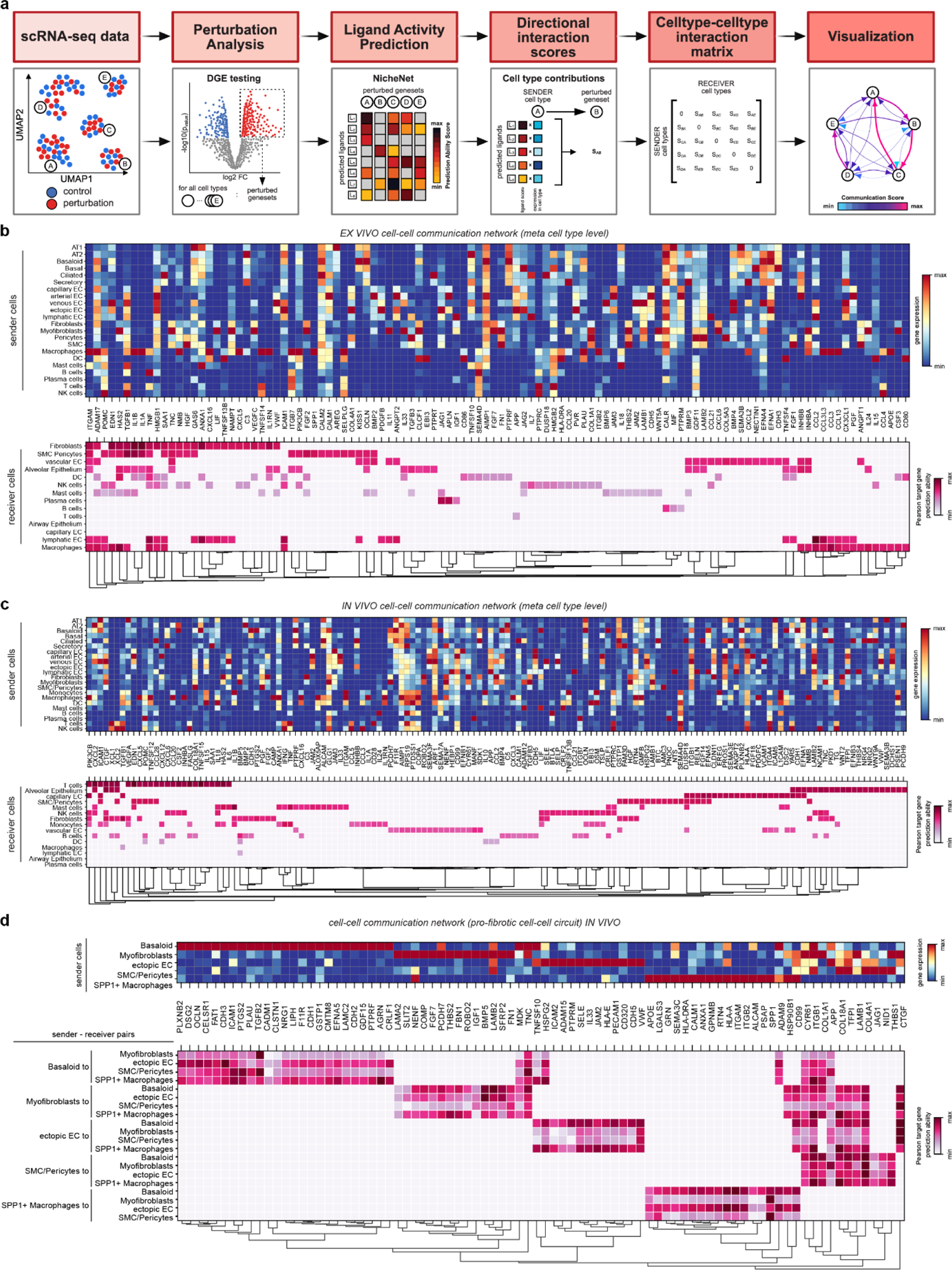
A conserved fibrogenic cell-cell communication circuit in hPCLS (corresponding to Figure 7). **(a)** Schematic illustration of the cell-cell communication analysis. Differential gene expression (DGE) testing was performed for each cell type to define perturbed gene programs. For each receiver cell type the respective perturbed gene programs were then used as input for NicheNet to predict ligands that could induce the gene program in the receiver cell type. For each sender-receiver cell type pair we then calculated the sum of the products of the ligand activity scores and the average expression of the ligand in a sender cell type to get a measure of how much a sender cell type contributes to the perturbation gene program in the receiver cell type. These so-called directional interaction scores in their entirety allowed the generation of a cell type-cell type interaction matrix, which was used to quantitatively visualize the directionality of cell-cell communication routes after perturbation. **(b,c)** Overview of the cell-cell communication routes between all meta cell states as predicted by NicheNet *ex vivo* **(b)** and *in vivo* **(c)**. The top panel shows the average expression of ligands predicted by NicheNet across sender cells. The bottom panel visualizes the corresponding predictive score of all ligands to induce their downstream target gene signature in the respective receiver cell types as measured by the Pearson correlation target gene prediction ability. Only top 25 ligands per receiver cell type are shown due to space constraints, which explains absence of some conserved ligands shown in **Fig. 7d**. **(d)** Overview of the cell-cell communication routes between five cell states of the fibrogenic alveolar cell circuit *in vivo* as predicted by NicheNet. The top panel shows the average expression of ligands predicted by NicheNet across sender cells. The bottom panel visualizes the corresponding predictive score of all ligands to induce their downstream target gene signature for the respective sender-receiver cell type combination as measured by the Pearson correlation target gene prediction ability.

**Supplementary Figure S12:**
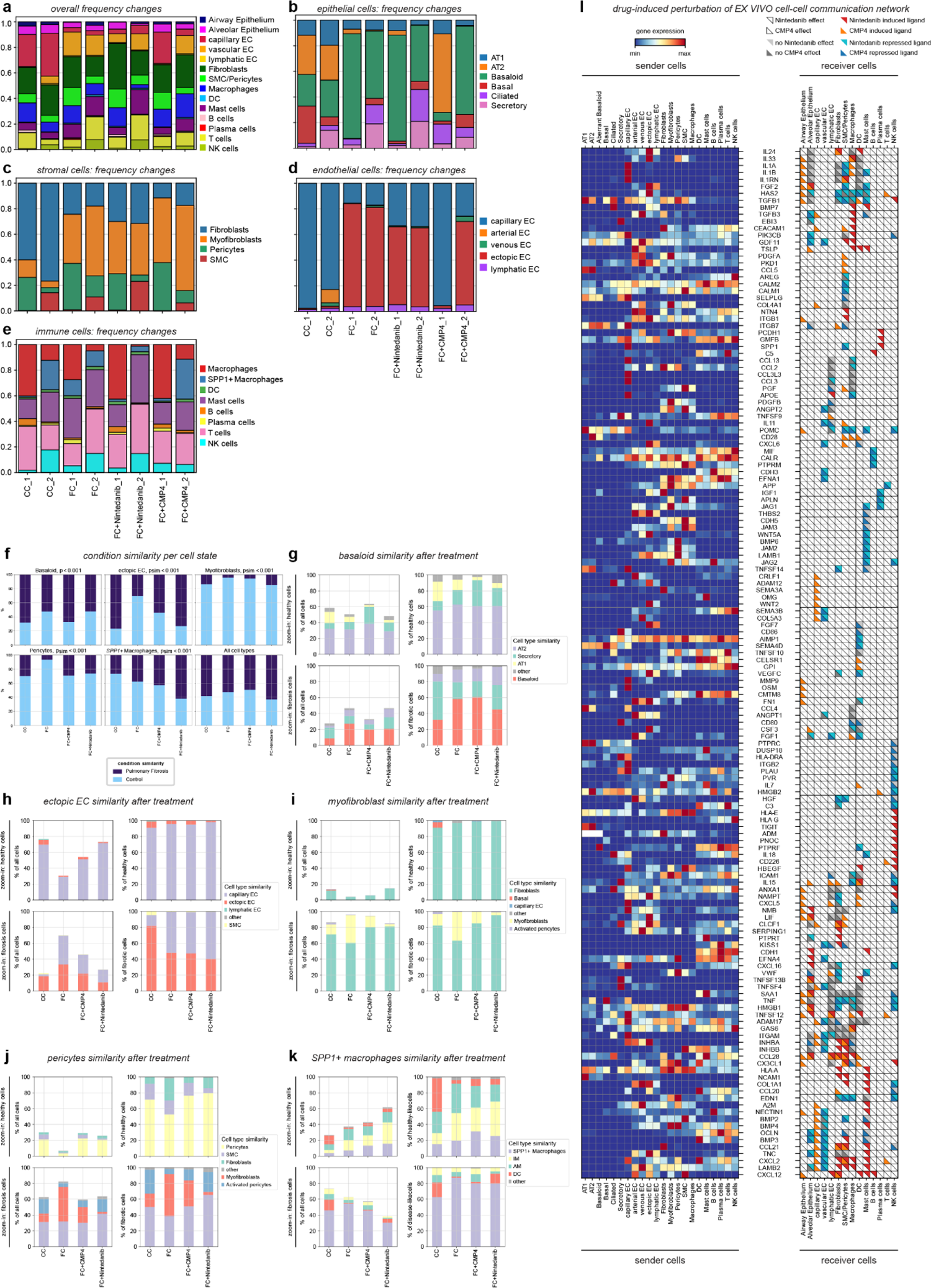
Drug-specific perturbations of the conserved cell-cell circuit (corresponding to Figure 8). **(a-e)** Cell type frequencies after CC, FC, FC+CMP4, and FC+Nintedanib treatment. **(f)** Similarity of CC, FC, FC+CMP4, and FC+Nintedanib treated cells from hPCLS compared to Control and Pulmonary Fibrosis cells in the PF-extended HLCA (*in vivo* reference) as assessed by scArches mapping. Stacked bar plots show the percentage of cells mapping to either Control or Pulmonary Fibrosis with regards to treatment for each cell state. Fisher’s exact test. **(g-k)** Shifts in cell type similarity of basaloid cells **(g)**, ectopic ECs **(h)**, myofibroblasts **(i)**, activated pericytes **(j)**, and SPP1+ macrophages **(k)** from CC, FC, FC+CMP4, and FC+Nintedanib treated hPCLS as assessed by scArches mapping. Upper rows show zoom-in on cells from healthy control lungs, lower rows show zoom-in on cells from pulmonary fibrosis lungs. **(l)** Heatmap visualizing perturbations of cell-cell communication routes after drug treatments as shown in Fig. 8h but based on all meta-cell types contained in the data set.

